# High throughput peptidomics elucidates immunoregulatory functions of plant thimet oligopeptidase-directed proteostasis

**DOI:** 10.1101/2022.05.11.491536

**Authors:** Anthony A. Iannetta, Philip Berg, Najmeh Nejat, Amanda L. Smythers, Rezwana R. Setu, Uyen Wesser, Ashleigh L. Purvis, Zoe A. Brown, Andrew J. Wommack, Sorina C. Popescu, Leslie M. Hicks, George V. Popescu

**Author notes:** These authors contributed equally to this work (Anthony A. Iannetta, Philip Berg). **Corresponding authors:** Dr. Leslie M. Hicks, Department of Chemistry, the University of North Carolina at Chapel Hill, Kenan Laboratories, 125 South Road, CB#3290, Chapel Hill, NC 27599-3290, United States, Dr. George V Popescu, Institute for Genomics, Biocomputing, and Biotechnology, Mississippi, State University, Mississippi State, MS, USA.

## Abstract

Targeted proteolysis activities activated during the plant immune response catalyze the synthesis of stable endogenous peptides. Little is known about their biogenesis and biological roles. Herein, we characterize an *Arabidopsis thaliana* mutant *top1top2* in which targeted proteolysis of immune-active peptides is drastically impaired during effector-triggered immunity (ETI). For effective ETI, the redox-sensitive thimet oligopeptidases TOP1 and TOP2 are required. Quantitative mass spectrometry-based peptidomics allowed differential peptidome profiling of wild type (WT) and *top1top2* mutant at the early ETI stages. Biological processes of energy-producing and redox homeostasis were enriched, and *TOPs* were necessary to maintain the dynamics of ATP and NADP(H) accumulation in the plant during ETI. Subsequently, a set of novel TOPs substrates validated *in vitro* enabled the definition of the TOP-specific cleavage motif and informed an *in-silico* model of TOP proteolysis to generate bioactive peptide candidates. Several candidates, including those derived from proteins associated with redox metabolism, were confirmed *in planta*. The *top1top2* background rescued WT’s ETI deficiency caused by treatment with peptides derived from targeted proteolysis of the negative immune regulator FBR12, the reductive enzyme APX1, the isoprenoid pathway enzyme DXR, and ATP-subunit β. These results demonstrate TOPs role in orchestrating the production and degradation of phytocytokines.

## Introduction

Plants have developed an innate immune response with complex, chemical-based signaling pathways to sense and respond to unfavorable environments (Chagas et al., 2018). Effector-triggered immunity (ETI) is a robust resistance response to virulence effectors deployed by pathogens to suppress and interfere with pathogen-associated molecular pattern-triggered immunity (Jones and Dangl, 2006). ETI is activated when nucleotide-binding leucine-rich repeat immune receptors recognize effectors, such as avrRpt2 from *Pseudomonas syringae* (Cui et al., 2015; Axtell and Staskawicz, 2003; Jones and Dangl, 2006). The detection of these effectors elicits the hypersensitive response (HR), a form of programmed cell death (PCD) that restricts pathogen growth (Lam et al., 2001; Heath, 2000). ETI execution is facilitated by a rapid increase in production and subsequent accumulation of reactive oxygen species (ROS), an event termed the ‘oxidative burst,’ leading to extensive post-translational protein thiol oxidation of the ETI proteome (McConnell et al., 2019). ROS accumulation elicited by ETI is biphasic with a low amplitude transient first phase, followed by a sustained phase of much higher magnitude (Torres et al., 2006; Lamb and Dixon, 1997). A positive feedback loop between ROS and defense hormones is maintained until a threshold is reached to trigger immune signal propagation, leading to ETI transcriptional reprogramming (Yoshimoto et al., 2009; Zurbriggen et al., 2010). During ETI and following the pathogen-triggered oxidative burst, damaged proteins with irreversibly oxidized residues accumulate at the site of pathogen infection; their timely removal is crucial to maintaining proteostasis (Das and Roychoudhury, 2014; Bassham, 2007).

Proteolysis is an irreversible protein post-translational modification essential for the functional regulation of numerous physiological and pathological processes (Rawlings and Salvesen, 2013). Complete proteolysis contributes to the maintenance of cellular proteostasis through protein degradation and turnover (Van der Hoorn and Klemenčič, 2021). In contrast, targeted or limited proteolytic cleavage can generate stable protein fragments with biological activity. More recently, targeted proteolysis has been recognized as a significant regulatory process of organismal response to pathogen attacks and developmental cues (Segonzac and Monaghan, 2019; Chen et al., 2020; Wang et al., 2022). A rich repertoire of plant proteases and peptidases, estimated to represent approximately 3% of the plant genome (Van der Hoorn, 2008; Paulus and Van der Hoorn, 2019), catalyzes degradation via peptide bond hydrolysis. Proteolysis produces rapid and substantial changes in protein dynamics of biological systems through the activation of highly regulated proteolytic cascades (Cheng et al., 2015; Paulus et al., 2020; Paulus and Van der Hoorn, 2019). Cytosolic proteolytic cascades complete the processing of proteasome-released peptides, whereas organelle-localized proteolytic components (*e.g.,* chloroplast and mitochondria) have a wide range of functions, including the processing of signal peptides during organellar import and removal of damaged proteins (Kidrič et al., 2014; van Wijk, 2015). Although connectivity between organellar and cytosolic proteolytic networks is not well characterized in plants, it is an essential component of the metazoan response to oxidative stress and pathogen attack (Suhm et al., 2018; Samant et al., 2018; Díaz-Villanueva et al., 2015). Substrates for diverse proteolytic cleavage have been identified in many plant species (Ziemann et al., 2018; Stegmann et al., 2017; Cheng et al., 2015). Nevertheless, the repertoire of plant peptides and functional roles of bioactive peptide products and their biogenesis remain largely uncharacterized.

Metazoan oligopeptidases with specificity limited to a few substrates are critical in generating bioactive peptides for stress response signaling through controlled proteolysis (Kessler et al., 2011; Ferro et al., 2014). Although these proteolytic processes are less explored in plants, plant peptidases have demonstrated other functions besides their role in protein homeostasis maintenance, including the release of defense response peptides (Tavormina et al., 2015). Endogenous peptides, termed phytocytokines, are primarily produced following partial proteolytic cleavage of precursor proteins in response to pathogen infection and amplify immune signals as part of feed-forward cellular circuits. Controlled proteolysis triggers ETI activation through specific receptor-effector recognition events. For example, the Arabidopsis immune receptor RPS2 senses the *Pseudomonas syringae* effector protease AvrRpt2 via cleavage of the guardee RIN4, prompting ETI activation (Axtell and Staskawicz, 2003); the *P. syringae* protease AvrPphB then cleaves the kinase PBS1 guarded by the immune receptor RPS5, initiating the recognition of PBS1 by RPS5 and ETI activation (Shao et al., 2003; Qi et al., 2014). These observations suggest a critical regulatory role of plant peptides in immune signaling.

TOP1 and TOP2 are zinc-dependent peptide hydrolases (Kmiec et al., 2016; Gomis-Rüth, 2009). These metallopeptidases are critical components in plant response to oxidative stress through SA-mediated signaling pathways and are required for a fully functioning immune response to ETI-activating pathogens (Moreau et al., 2013; Westlake et al., 2015). The Arabidopsis genome contains three genes encoding TOPs, two of which have been characterized in depth: *TOP1* and *TOP2*. TOP1 (AT5G65620, also named organellar oligopeptidase, OOP) contains an N-terminal signal peptide that mediates its localization to the chloroplast and mitochondria (Kmiec et al., 2013; Moreau et al., 2013). TOP1 cleaves presequences containing 8-23 amino acids *in vitro* and is hypothesized to act downstream of organellar proteases for intra-organelle peptide degradation and organelle import processing (Kmiec et al., 2013). TOP2 (AT5G10540, also known as cytosolic oligopeptidase, CyOP) functions downstream of the 20S proteasome, degrading proteasome-generated peptides during oxidative stress (Polge et al., 2009; Moreau et al., 2013; Kmiec et al., 2013). Prior evidence suggests that TOP1 and TOP2 have functional overlap in ETI and PCD (Westlake et al., 2015; Polge et al., 2009; Kmiec et al., 2013; Moreau et al., 2013). Both oligopeptidases are required for plant defense against avirulent strains of *P. syringae* through the activation of the resistance proteins RPS2 or RPS4 and both are necessary to regulate PCD (Moreau et al., 2013). Indirect evidence supports a role for TOPs in the controlled proteolysis of rotamase cyclophilin 1 (ROC1/CYP18–3), required for AvrRpt2 protease self-cleavage prior to the activation of ETI (Al-Mohanna et al., 2021). In a current model, TOPs are components in an interconnected organelle and cytosol proteolytic pathway that regulates the ETI oxidative burst and pathogen resistance through SA, ROS, and antioxidants (Westlake et al., 2015).

When nullified via genetic or chemical approaches, peptidase deficiency leads to a decrease in the accumulation of products and an increase in substrates (Lone et al., 2013; Cavalcanti et al., 2014). We hypothesized that the absence of TOP1 and TOP2 would increase the intracellular abundance of TOP peptide substrates, as evidenced when comparing *top1top2* with WT; therefore, peptides over-accumulating in the *top1top2* may represent direct or indirect TOPs substrates. Likewise, a significant increase in the quantity of products derived from these substrates would be expected in WT compared to the mutant. We used the double mutant instead of *top1* and *top2* single mutants for comparative analysis due to their documented shared roles in ETI and PCD (Westlake et al., 2015; Polge et al., 2009; Kmiec et al., 2013; Moreau et al., 2013). Furthermore, while TOP1 and TOP2 have different subcellular localizations, their functional overlap and high sequence similarity suggest a potential for redundant proteolytic activity and substrates (Kmiec et al., 2016; Moreau et al., 2013).

Our prior work delineated TOP peptide substrates via quantitative *in vivo* peptidomics comparing Arabidopsis WT and the *top1top2* (Iannetta et al., 2021). Herein, we implemented a similar approach to characterize the peptidomes of pathogen-infected *A. thaliana* wild type (WT; Col-0 accession) and *top1top2* plants during the early stages of ETI. Our characterization revealed the ETI peptidome and its temporal dynamics at critical time points post-infection. Differential peptide analysis generated a set of potential TOP substrates and an *in-silico* model of TOPs proteolytic activity. A search for bioactive peptides associated with TOP activity yielded candidates with unique sequence characteristics, whose roles in ETI were tested in WT and *top1top2*. These results highlight the complex dynamics of proteolytic events during the plant immune response. We show that predicted peptides can strongly modulate ETI phenotype in both genotypes and successfully rescue the ETI defective phenotype of *top1top2.* We propose that TOPs are a powerful model for studying the coordination between controlled proteolysis and immunity in plants.

## Results

### The ETI-triggered peptidome

To measure peptidome changes during the initial stages of ETI and elucidate TOP-mediated proteolytic pathways in plant defense, WT and *top1top2* rosette leaves were analyzed at 0 minutes post-inoculation (mpi), 30 mpi, and 180 mpi after inoculation with *Pseudomonas syringae* pv. tomato DC3000 carrying the avirulence gene avrRpt2 (*Pst*AvrRpt2). Timepoint selection was designed to capture differential proteolytic events occurring in the early stages of the ETI oxidative burst and the altered dynamics of the *top1top2* delayed activation of ETI (McConnell et al., 2019).

Quantitative peptidomics was performed on three biological replicates at each time point in both genotypes (**Figure 1**). The analysis of the WT samples produced 2810 quantifiable peptides from 698 proteins, while *top1top2* revealed 2793 quantifiable peptides from 693 proteins (**Supplemental Data Set S1**).

**Figure 1:**
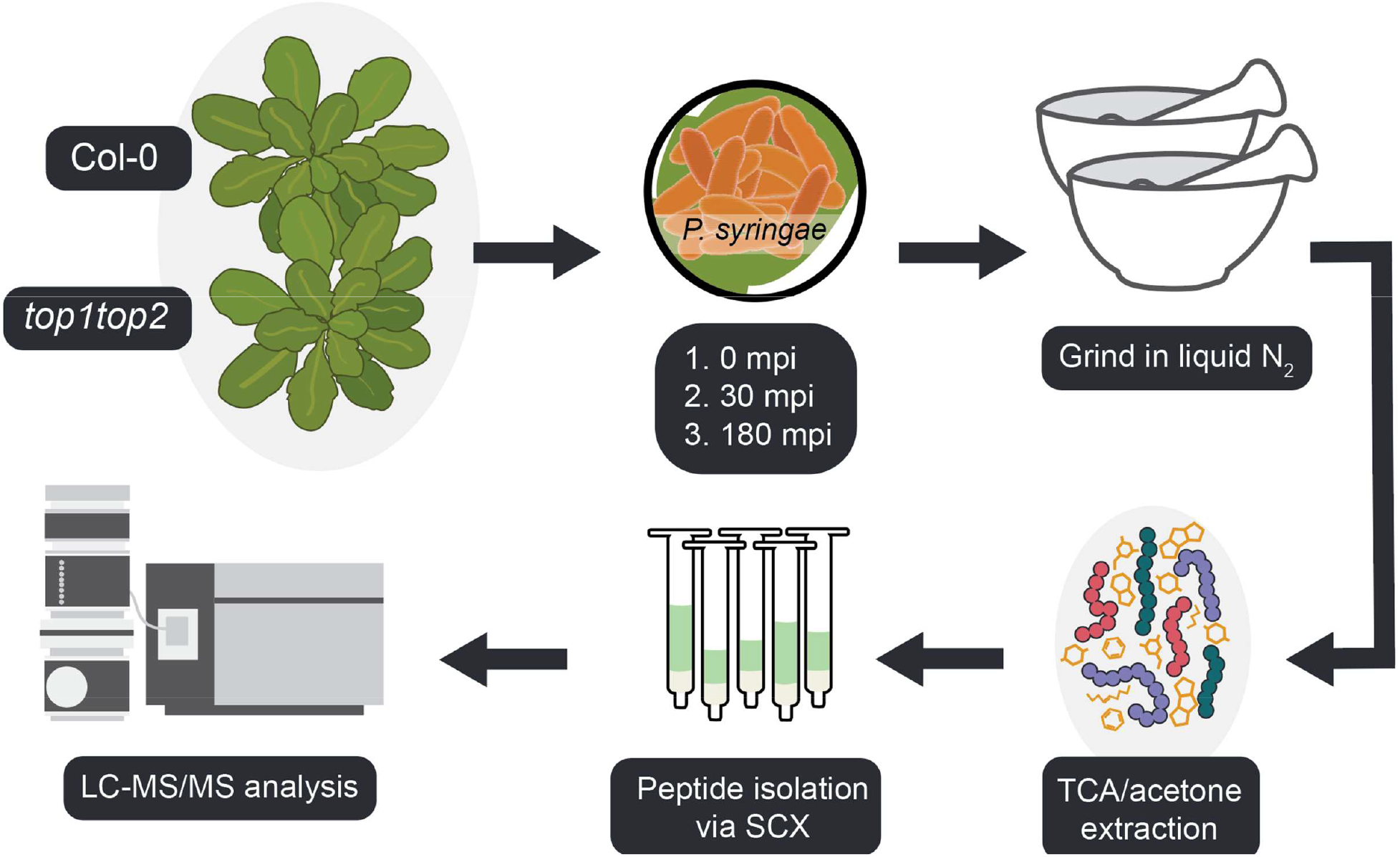
The peptidomes of WT and *top1top2* rosette leaves were analyzed following pathogen inoculation with *Pst* avrRpt2 to measure peptidome changes during the initial stages of ETI and elucidate TOP-mediated proteolytic pathways during plant defense. Rosette leaves were ground under liquid nitrogen before extracting peptides with 10% TCA in acetone. Peptides were isolated from small molecules with SCX SPE before peptide quantitation. Peptide concentrations across replicates were quantified and normalized before liquid chromatography-tandem mass spectrometry (LC-MS/MS) analysis.

### Differential peptidomics reveals potential TOP substrates during ETI

Two approaches were taken to assess peptide abundance differences across these conditions. First, peptide abundances were compared across infection time points within each genotype to determine peptidome differences during defense response. Second, peptide abundances were compared across genotypes to characterize ETI-mediated TOP proteolysis. Differentially abundant peptides (DAPs) between the WT and *top1top2* genotypes were identified using our pipeline (Berg et al., 2019) for label-free quantification of post-translational modifications (PTMs), as described in the methods section. Overall, 325 peptides significantly accumulated in WT and 125 accumulated in *top1top2* at 0 mpi. At 30 mpi, 276 peptides accumulated in WT and 246 accumulated in *top1top2*, while at 180 mpi 59 peptides accumulated in WT and 64 accumulated in *top1top2* (**Figure 2A, Supplemental Data Set S2**). The analysis of DAPs revealed that most were unique to either control (0 mpi) or early (30 mpi) *Pst*AvrRpt2 infection (**Figure 2B**). There are significant differences between the peptidomes of WT and *top1top2* at 0 mpi, indicating compensatory effects (possibly transcriptional reprogramming) triggered by the absence of TOPs. The most significant differences between peptidomes, measured at 30 mpi, diminish at 180 mpi, suggesting convergence of the mutant and WT response dynamics in time. We also analyzed the temporal dynamics of the WT and *top1top2* peptidomes during ETI and found that peptide abundance fold-changes had a smaller range in *top1top2* series as compared to the WT series (**Supplemental Figure S1**). As a result, the *top1top2* differential peptidome was drastically reduced compared to the WT within time series comparisons (**Supplemental Figure S2, Supplemental Data Set S3**). Most DAPs were unique to one comparison indicating a high dynamic of proteolytic activity during ETI responses. The *top1top2* differential peptidome was more than three times smaller in WT at 180 mpi vs 30 mpi, indicating a significant deficiency in proteolytic activity associated with ETI.

**Figure 2:**
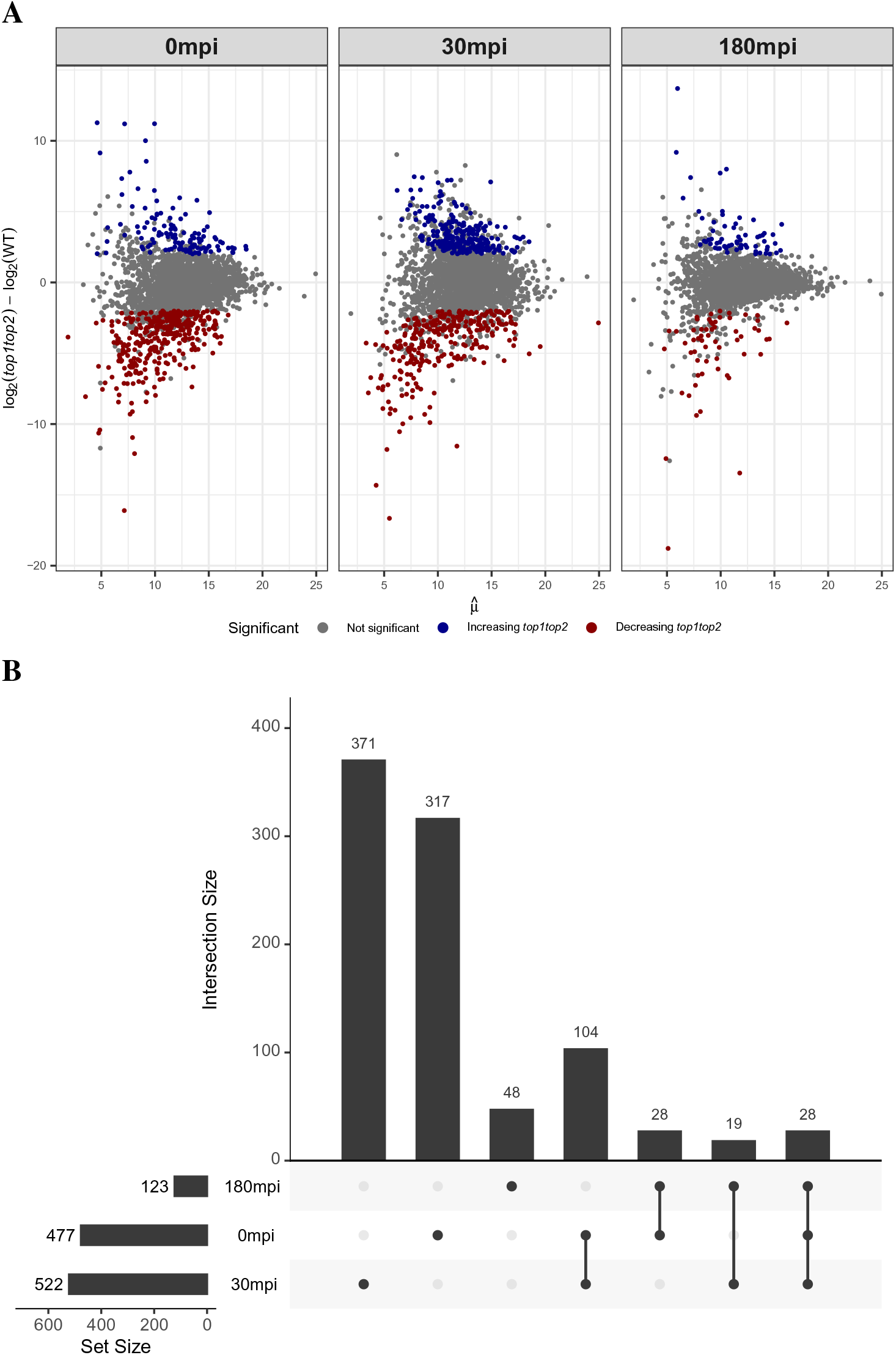
The distribution of the decision of the differential abundance analysis for the peptides. (**A**) shows a MA-plot with the title of each facet corresponding to the decision of that comparison. Each dot corresponds to a peptide, and the x-axis shows the mean of that peptide for that comparison, while the y-axis shows the log fold-change, and a positive value indicates accumulation in *top1top2*. Peptides that were imputed were represented with their median mean and fold change from all imputations. (**B**) UpSetR analysis of the significant hits. The figure shows the size of the set(s) marked with a dot below the bars.

### Functional characterization of the differential peptidome

Since peptides accumulated in either genotype represent potential TOP substrates or TOP-cleaved peptide products, the DAP comparisons can reveal TOP-mediated proteolytic activity pre- and post-inoculation with the pathogen. We performed a gene ontology (GO) term enrichment analysis of DAPs to identify the molecular function and biological processes associated with TOP-mediated proteolysis. GO enrichment analysis using *ThaleMine* (Krishnakumar et al., 2017) identified 18 common categories for the analyzed time points (**Supplemental Data Set S4**). Significant enriched GO terms across all time points included ‘metabolic processes’, ‘photosynthesis’, and ‘peptide biosynthetic processes’; other categories of interest were ‘translation’ and ‘response to metal ions’ (**Figure 3A**). A drastic change in unique GO terms was observed between 0 mpi (‘cytoplasmic translation’ and ‘electron transport chain’) and 30 mpi (22 terms including ‘regulatory metabolic processes’, ‘ATP production’, and ‘ribosome biogenesis’), indicating a significant change in the landscape of TOP proteolyzed proteins during ETI (**Figure 3B**). The GO analysis of the peptidome temporal dynamics identified common GO categories (**Supplemental Figure S3**) as well as additional unique GO terms that characterize WT and *top1top2* ETI responses (**Supplemental Figure S4**). Most of the categorical changes in GO biological processes occurred between 30 and 180 mpi for WT, whereas they occurred between 0 and 30 mpi in *top1top2*. The analysis shows a reduction in GO terms and overrepresentation levels in *top1top2* compared to the WT. Overall, these results validate TOPs proteolytic functions under physiological conditions (Westlake et al., 2015) and suggest an essential role in ETI’s temporal dynamics.

**Figure 3:**
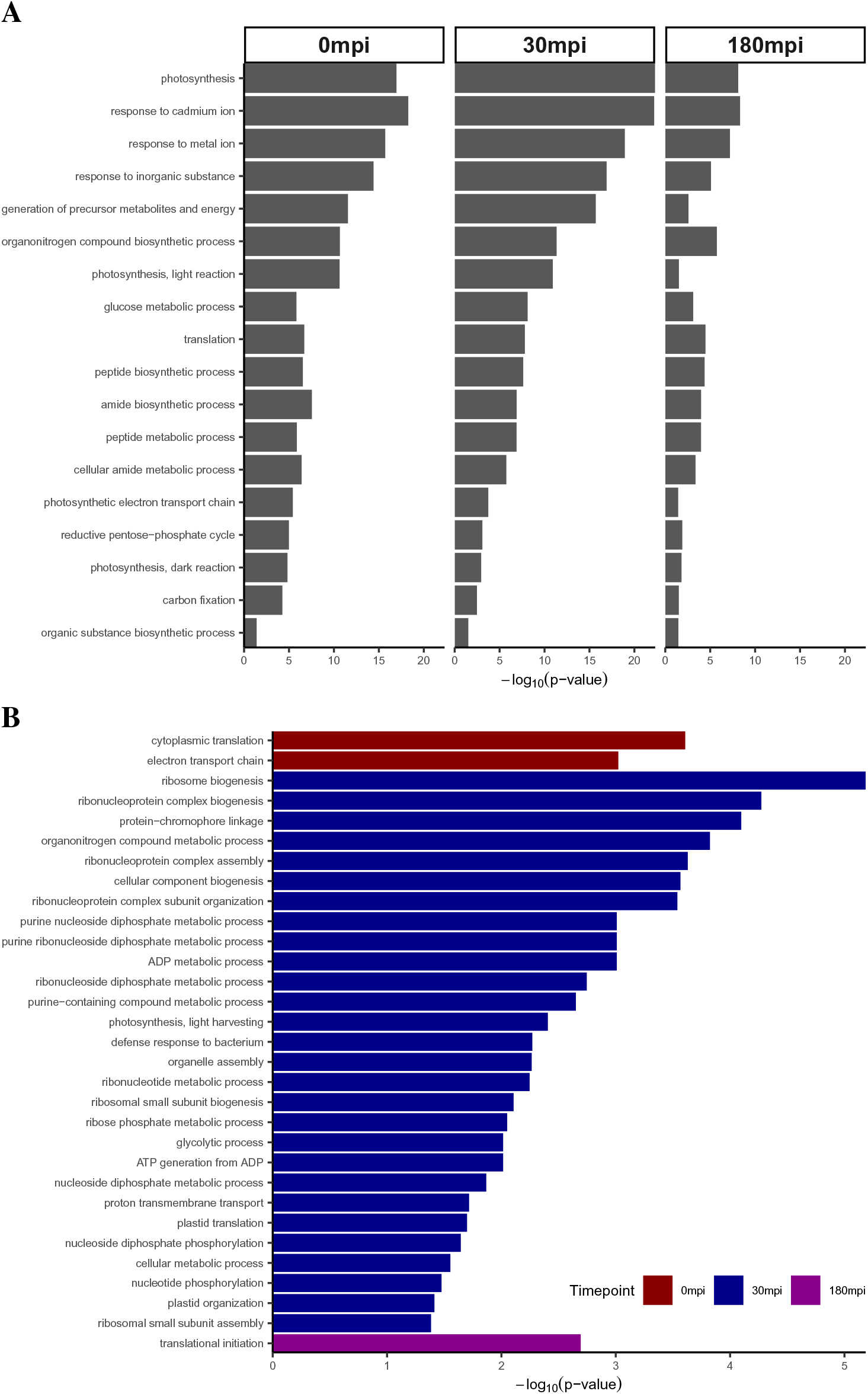
GO-enrichment of the differneitally abundant peptides at the different time points. The x-axis shows the category, and the y-axis shows the -log10(p-value) of the statistical test for overrepresentation. (**A**) shows common GO terms enriched in all time points with the title of the facets indicating the time point. (**B**) shows the GO terms unique to different time points. GO-enrichment of biological process unique to 0 mpi and 30 mpi for the significant proteins. Red color indicates the GO-terms unique to 0 mpi, and blue indicates the same but for 30 mpi.

### TOPs are required for metabolic homeostasis during the ETI immune response

Several peptides from chloroplast and mitochondrial ATP synthase subunits over-accumulated in *top1top2* relative to the WT and conversely decreased in abundance in WT post-inoculation with *Pst*AvrRpt2. GO analysis of DAPs also revealed enrichment for ATP generation as well as other ATP-related metabolic processes at 30 mpi. We hypothesized that TOPs are necessary for ATP synthase processing during the ETI; thus, in *top1top2,* ATP synthase proteostasis might be dysfunctional. To verify this hypothesis, ATP cellular concentration was measured in WT and *top1top2* plants in controls (0 mpi) and following inoculation with *Pst*AvrRpt2 at 30 and 180 mpi. The temporal dynamics of ATP concentration was markedly different between genotypes (**Figure 4A**). Whereas similar levels were recorded at 0 mpi, WT plants showed a significant burst in ATP accumulation at 30 mpi followed by a reversal to the 0 mpi level at 180 mpi. In contrast, *top1top2* plants lacked the 30 mpi burst, showing a slight increase over time in cellular ATP concentration.

**Figure 4:**
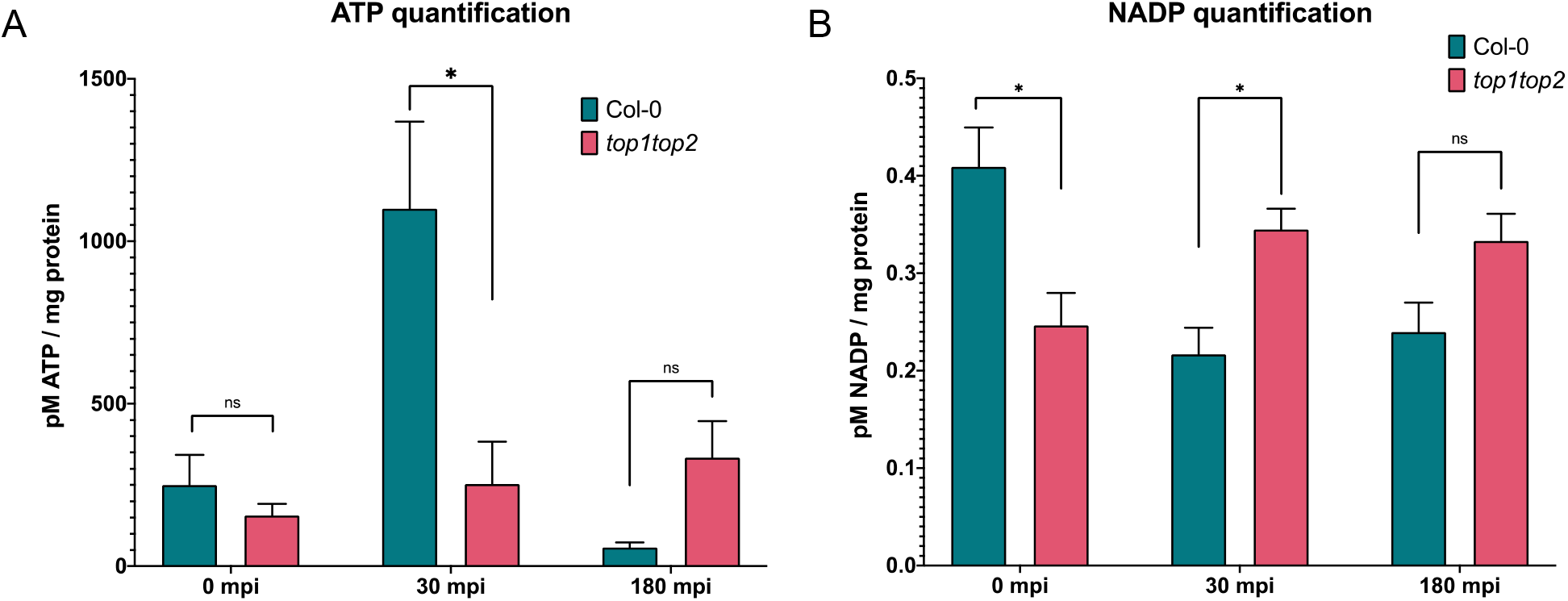
The metabolic state of Arabidopsis WT and *top1top2* plant lines following inoculation with *Pst* avrRpt2. (A) ATP was quantified using a luciferase assay (*: *p* < 0.05, ns: not significant, following two-sample equal variance *t*-test). (B) NADP^+^ and NADPH were quantified using an enzyme cycling assay (*: *p* < 0.05, ns: not significant, following two-sample equal variance *t*-test).

The ATP production burst at 30 mpi was coupled with perturbation in the dynamics of metabolic and energy processes, as evidenced by enrichment of the cognate GO categories at 0 mpi, 30 mpi, and 180 mpi (**Figure 3A**). In addition, the significant representation of “Reductive pentose-phosphate cycle” and “photosynthesis, dark reaction” GO categories in *top1top2* signify distinct dynamics of the NADP(H) reactions between genotypes. These observations point to a potential role for TOP-modulated proteostasis in this process, a hypothesis tested by measuring the cellular concentration of NADP(H) in both genotypes. NADP(H) concentration was measured in WT and *top1top2* at 0, 30, and 180 mpi following inoculation with *Pst*AvrRpt2 (**Figure 4B**). We observed a significantly higher initial (0 mpi) concentration of NADP(H) in WT compared to *top1top2*. NADP(H) cellular accumulation dynamics over the tested time interval were also markedly different between the genotypes. In WT, NADP(H) concentration decreased at 30 and 180 mpi, whereas in *top1top2,* NADP(H) concentration increased significantly at both 30 and 180 mpi. Such contrasting dynamics suggest a regulatory role for *TOPs* in cellular NADP(H) reduction during ETI.

### In vitro screening for TOP substrates

Pairwise comparisons between genotypes across infection time points identified peptides significantly more abundant in *top1top2* mutant plants. Peptides with increased abundance in the mutant likely represent either direct substrates or result from compensatory proteolytic activities due to the loss of TOPs (Rei Liao and van Wijk, 2019). Screening these peptides using *in vitro* peptidase assays helps differentiate between direct and indirect TOP substrates. For testing TOP substrate candidates, purified TOP1 and TOP2 were incubated with candidate peptides. We selected peptides for TOPs substrate screening using the following criteria: *1)* consideration of previously determined TOP substrate length (8-23 amino acids), *2)* detection of increased peptide abundance in *top1top2* over multiple time points, *3)* detection of increased abundance of peptides in *top1top2* from the same protein with overlapping recognition sequences and *4)* large *in vivo* fold changes in peptide abundance. The *in vitro* enzyme assays were conducted with the 13 synthetic peptides (**Supplemental Figure S5**) that met the above criteria. The selected peptides were derived from proteins involved in photosynthesis, ATP synthesis/binding, carbon fixation, and fatty acid synthesis/beta-oxidation. Substrate candidates were incubated with His-tagged recombinant isoforms of TOPs — a TOP1 isoform lacking the organellar signal peptide (Gomis-Rüth, 2009), herein named ChlTOP1, and TOP2 — followed by LC-MS analysis. Of the 13 candidate peptides, 10 were identified as cleaved by ChlTOP1 and/or TOP2, representing direct TOPs substrates (**Supplemental Table S1**); ChlTOP1 had more cleavage sites (10 confirmed substrates, 18 total cleavages) than TOP2 (8 confirmed substrates, 15 cleavage sites).

### Characterization of TOPs cleavage patterns using differential peptide analysis

To characterize patterns of proteolytic activity in our dataset, we first examined peptides that are significantly more abundant in WT than in *top1top2* across the analyzed time points. Motif analysis was performed on these prospective TOP recognition sites using PLogo (O’Shea et al., 2013) after extending peptide termini to include the surrounding amino acids (AA). The resulting cleavage pattern shows a significant enrichment of Val at P4, Ala at P3, Val and Pro at P2, Lys at P1, Ala at P1’, Val at P2’, Ala at P3’, and Asp at P4’ (**Supplemental Figure S6**), but no strong similarity with other reported M3 metallopeptidase cleavage patterns (Rawlings, 2016).

We next used the peptides from the *in vitro* screen described above and previously published substrates (Al-Mohanna et al., 2021; Iannetta et al., 2021) to learn TOPs cleavage patterns. Further, we considered all peptides with a –log_10_(p-value) <0.29 and a |LFC|<0.5 in all three time points as negative examples. We first analyzed the amino acid composition of the peptides with at least one validated cleavage site. The most common amino acids were Ala and Gly, both small and hydrophobic, while Cys and Trp were never observed (**Supplemental Figure S7**). Next, we looked at the amino acid composition around the cleavage sites. We found that Pro never occurred at P1’ but was in high frequency at P2’ (**Supplemental Figure S8**). We also observed that Ala tends to be found at a higher frequency toward the C-terminus. Toward the N-terminus, we observed a preference for Leu and Gly at P1 and Leu at P2. We also found a general preference for hydrophobic amino acids around the cleavage sites.

To increase pattern specificity, we next mined amino acid properties at predicted TOP cleavage sites. Hydrophobic (and amphipathic) amino acids were classified into three categories: *a)* large (l), consisting of Trp, Phe, and Tyr, *b)* small (s) consisting of Gly and Val, and *c)* medium (o) containing Ile, Leu, Val, and Met. Due to its distinctive pattern at P2’, Pro was assigned its own category (p). Asp and Glu were classified as negatively charged (n), while Arg, Lys, and His were classified as positively charged (r). Ser, Thr, Cys, Gln, and Asn were classified as hydrophilic (i). We discovered a general preference for small amino acids toward the C-terminus. At the same time, there was a strong preference for hydrophobic amino acids toward the N-terminus, especially at P2, and that large amino acid had low frequency toward both termini and a weak preference for negatively charged amino acids at P1 and P1’ (**Supplemental Figure S9**).

Next we assessed cleavage motif uncertainty to understand putative interactions between pairs of amino acids at the cleavage site. We calculated the Shannon entropy and the mutual information (Thomas and Joy, 2006) in an eight amino acid window (P4 to P4’) centered at the cleavage sites. For peptides that were not cleaved, we calculated these statistics by sliding windows of eight amino acids across all detected peptides lacking a validated cleavage site (**Figure 5**). Shannon entropy (the antidiagonal) can highlight sites with a higher degree of uncertainty. At the same time, mutual information can help identify patterns of AA interaction (i.e., knowing the distribution of AA in one position informs about the distribution of AA in another position). We found that the entropy was reduced at most cleaved vs. uncleaved peptides positions, reaching the smallest values at P2, P2’ and P1’. At the same time, there was a substantial increase in mutual information (up to ten-fold) between amino acid positions in cleaved vs. the uncleaved peptides. Further, P3’ and P1 tended to have a larger mutual information with other cleavage site positions while P2’ shared less information with the other positions. Taken together, this indicates that the patterns of amino acid properties determining TOPs cleavage recognition are more specific than the general peptidome cleavage motif and could be used for substrate prediction.

**Figure 5:**
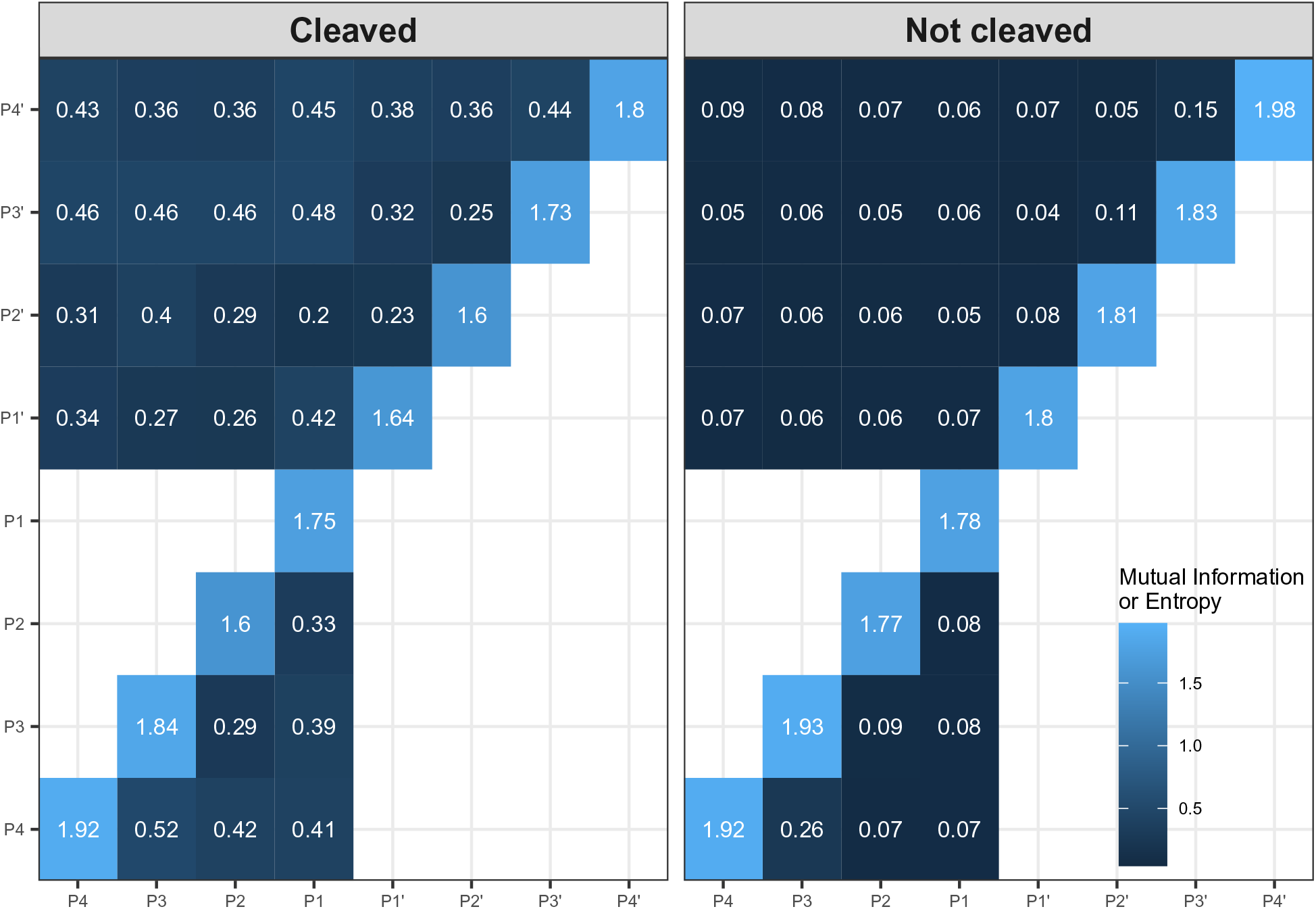
Comparison of the information of the amino acid properties around the validated cleavage sites and for a sliding window of size six for the amino acid properties in the peptides not cleaved as well as the negative examples. The x- and y-axes show the different positions around either the cleavage sites or the sliding window. The diagonal shows the entropy values, and the remaining cell show the mutual information between the positions.

### A predictive model of TOP proteolytic activity

To generate a predictive cleavage pattern of TOPs, we used a least-squares support vector machine (SVM) with a Laplace kernel (Suykens and Vandewalle, 1999). SVMs have been shown to have good performance and avoid overfitting when predicting peptide cleavage sites (duVerle and Mamitsuka, 2012). The model was trained on the *in vitro* validation data and used to predict cleavage sites in the entire stress peptidome (extending sequences with up to three amino acids from the protein sequence to detect cleavage sites at peptide termini). We analyzed 47,454 windows in the entire peptidome, from which we predicted 405 putative TOPs cleavage sites (**Supplemental Data Set S5**). Out of these, 173 cleavage sites mapped to peptides with significantly different abundance between *top1top2* and WT were selected for analysis. Of these, 65 cleavage sites were coherent with the observed peptide abundance fold-changes and could be associated with TOPs proteolytic activity: that is, if the cleavage was predicted in the observed peptide it accumulated in *top1top2*, while if predicted in the termini it accumulated in Col-0.

Peptides containing these putative TOPs cleavage sites were divided into two categories: a) TOPs substrate peptides (TSP) - the cleavage site was predicted in the observed peptide, and it accumulated in *top1top2*, b) TOPs product peptides (TPP) - the cleavage site was predicted in the peptide termini, and it accumulated in WT. Out of the 66 predictions, 54 were novel, and 12 were part of the training data (**Supplemental Table S2**). We then performed functional enrichment using *ThaleMine* (Krishnakumar et al., 2017) for the proteins corresponding to the TSPs and TPPs. The glucose metabolic process was enriched (**Figure 6**) mainly due to peptides originating from glyceraldehyde 3-phosphate dehydrogenases GAPA-2, GAPC2, and GAPB. Additionally, we observed TSPs from the ATP synthase delta subunit, possibly linking TOPs proteolytic activity to the observed ATP deficiency in the *top1top2* mutant. Analysis of logarithmic fold change (LFC) of TPPs and TSPs showed that the peptide with the largest LFC, a TPP, belonged to ribosomal protein RPS10C, while the third-largest LFC belonged to a GAPA-2, a TSP peptide. Other notable hits were a TSP belonging to chloroplastic DXR from the plastid non-mevalonate pathway, a TSP from the beta subunit of ATP synthase, and two peptides (a TPS and a TPP) from the L-ascorbate peroxidase 1 (APX1). Taken together, the model predicted TOPs proteolytic activity associated with peptides found in redox regulation and metabolic processes and provided possible links to the observed ATP synthesis molecular phenotypes.

**Figure 6:**
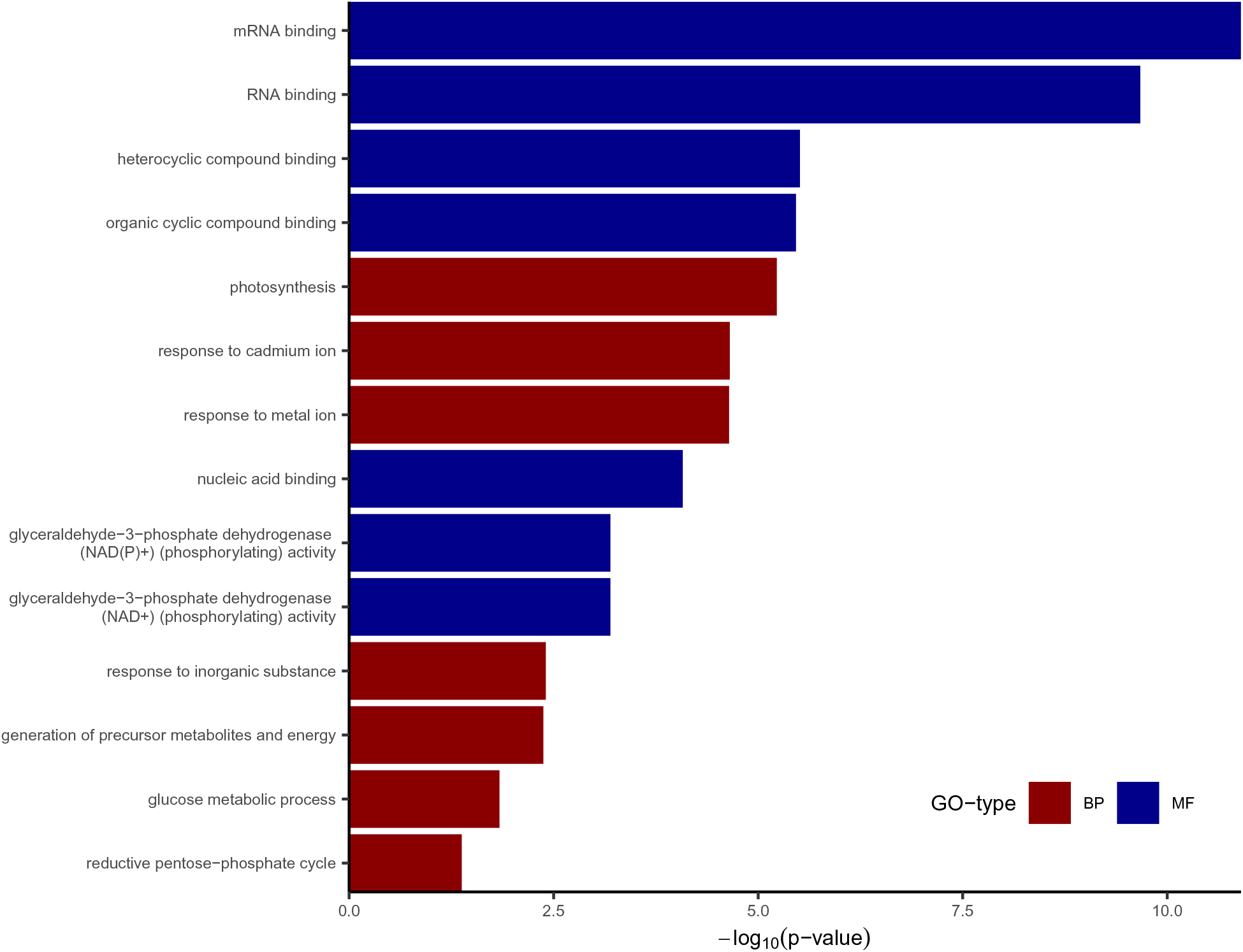
Shows the GO-enrichment of the proteins with peptides that had predicted cleavage sites that coincided with their fold-change. The labels correspond to the GO-term; red labels show biological processes while blue shows molecular function. The x-axis indicates the number of genes in the category, and the y-axis shows the -log10(p-value) of the statistical test for overrepresentation.

### TOPs cleavage motif analysis

We used pLogo (O’Shea et al., 2013) for statistical analysis and visualization of TOPs cleavage motif using the AA property classes described above. We contrasted cleavage motif characteristics of peptidome-wide predicted cleavagee sites vs. the TPP and TSP cleavage sites. In the peptidome-wide motif, the strongest preference was for proline at the P2’ position, and small amino acids were preferred at the P1, P1’, and P3’ positions (**Supplemental Figure S10**). In addition, negatively charged amino acids were lacking from all positions, while some hydrophilicity was preferred at P3 and P2. The TSP and TPP cleavage site motif strongly preferred positively charged and hydrophobic amino acids at P3 and large amino acids at P2 (**Figure 7A**). We also discovered an increased preference for negative amino acids at P1. The C-terminus motif showed small changes, with a strong selection for Pro at P2’, small amino acid at P1’, and positively charged amino acid preference at P3’.

**Figure 7:**
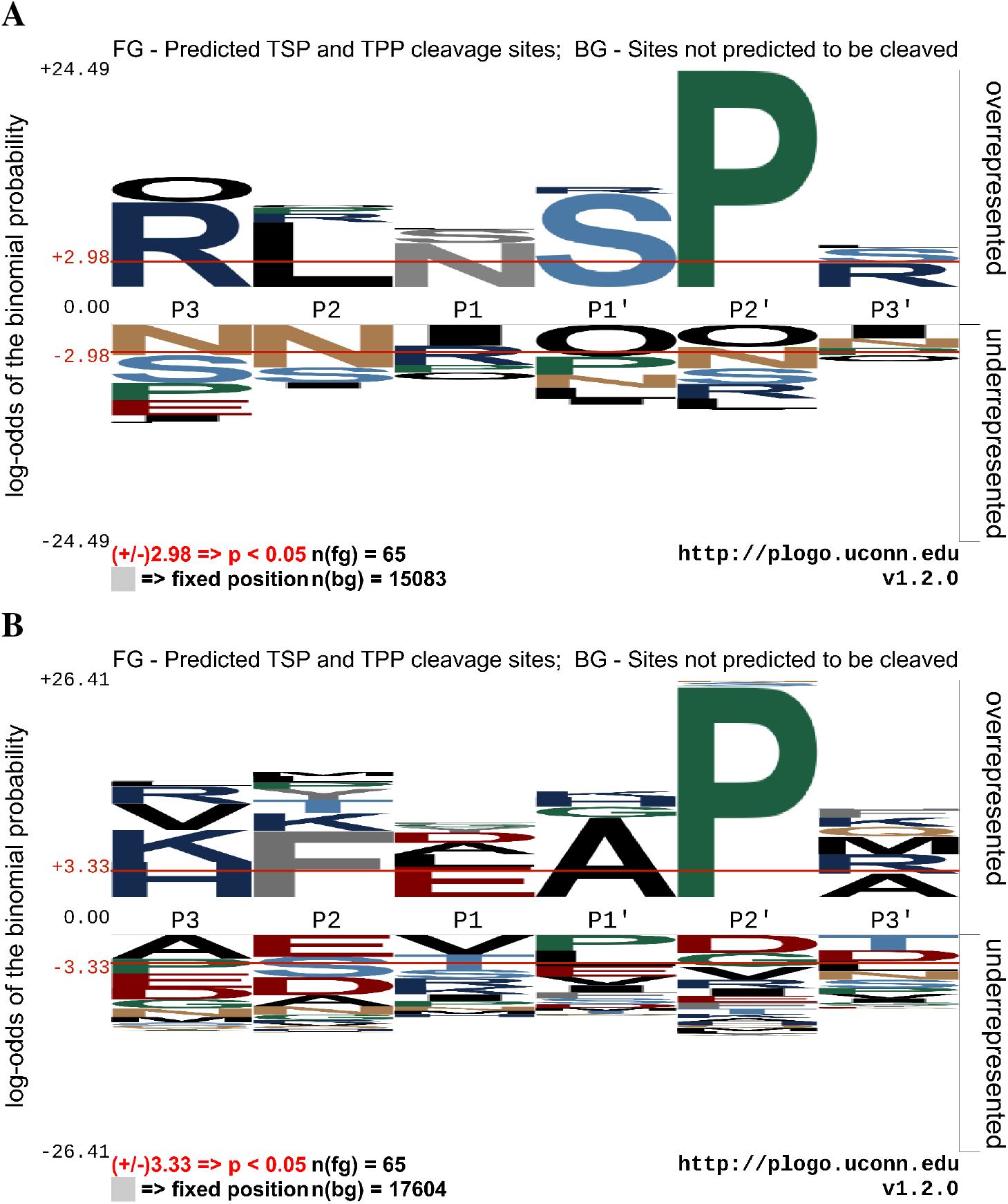
pLogo shows the overrepresentation of amino acid properties for the putative TOPs cleavage sites. (**A**) shows the motif for all sites predicted as cleaved. (**B**) shows the motif of the TPP and TPS.

We also analyzed the AA composition at predicted TPP and TSP cleavage sites (**Figure 7B).** Similarly, the primary signature of the cleavage motif was a strong preference for Pro at P2’, while P2 exhibited a strong preference for phenylalanine and P1’ for alanine. In addition, the peptidome-wide motif showed a preference for Ala at P1’ and P3’ (**Supplemental Figure S11**). The motifs analyzed here showed strong preferences for Pro and Phe, two key amino acids overrepresented in the proteolytic motifs of M3 metallopeptidases (Rawlings, 2016). However, the positions of the two signature amino acids were P2’ and P2 rather than P1’ and P1, while the motif specificity was lower, indicating a significant divergence of our peptidome analysis predictions from previously published MEROPS results.

### Prediction of bioactive peptides from differential peptidome

Another objective of our differential peptidome analysis was the identification of bioactive peptides with ETI-related signaling functions. To develop a bioactive peptide prediction model, we scored the peptide sequences using a Markov Chain (MC) describing amino acid transition probabilities (see Methods section). The model attempted to discriminate peptides with a “unique” primary sequence characterized by multiple low amino acids transition frequencies. We used a right-tailed Z-test to identify peptide sequences significantly different from the rest of the peptidome. To validate the model, we tested it on previously identified bioactive peptides involved in Arabidopsis stress signaling (Chen et al., 2020) and found that more than half had significantly high scores (**Supplemental Table S3**). We then applied the model to select a list of high-scoring peptides from our ETI-triggered peptidome; out of these, we selected 60 peptides (mapped to 35 proteins) with significantly high Z-scores and designated as DAPs (**Supplemental Table S4**).

GO enrichment analysis of these 35 proteins showed an over-representation of molecular functions related to translational regulation and mRNA binding (**Figure 8**). Four proteins participate in translation initiation: EIF4A1 (F4JEL5), EIF4A-2 (F4HV96), TRIP-1 (Q38884), and FBR12/eIF5A-2 (Q93VP3), and two proteins in translation elongation: TUFA (P17745) and eEF-1A1 (P0DH99). Most of these peptides were TOP substrates, except for two peptides from EIF4A1, which were putative TOP products. Among the predictions were two short proteins (100-140 amino acids) with disordered regions: AT2G33845 (Q8RYC3) and AT1G26550 (Q9FE18) encoded by mobile RNAs moving between various organs under normal or nutrient-limiting conditions (Thieme et al., 2015). We also found several GO enrichment categories related to stress signaling containing targets from redox-related proteins. Two of these peptides, AEQAHGANSGIHIA and GPDIPFHPG, mapped to APX1 (Q05431), an antioxidant enzyme with a role in redox homeostasis, while a third peptide, ASIKVHGVPMSTA, mapped to GSTF8 (Q96266), a chloroplast glutathione *S*-transferase. Another intriguing bioactive peptide candidate, AMKDAIEGMNGQDLDGR, mapped to GRP7 (Q0325), a small RNA binding protein that is a part of a circadian clock-regulated toggle switch involved in stomatal opening control (Schmal et al., 2013). Lastly, we predicted two bioactive peptide candidates in ATP-synthase subunits alpha (P56757) and two in the beta subunit (P19366). All four peptides were putative TOP substrates and may be associated with the ATP molecular phenotype of *top1top2*.

**Figure 8:**
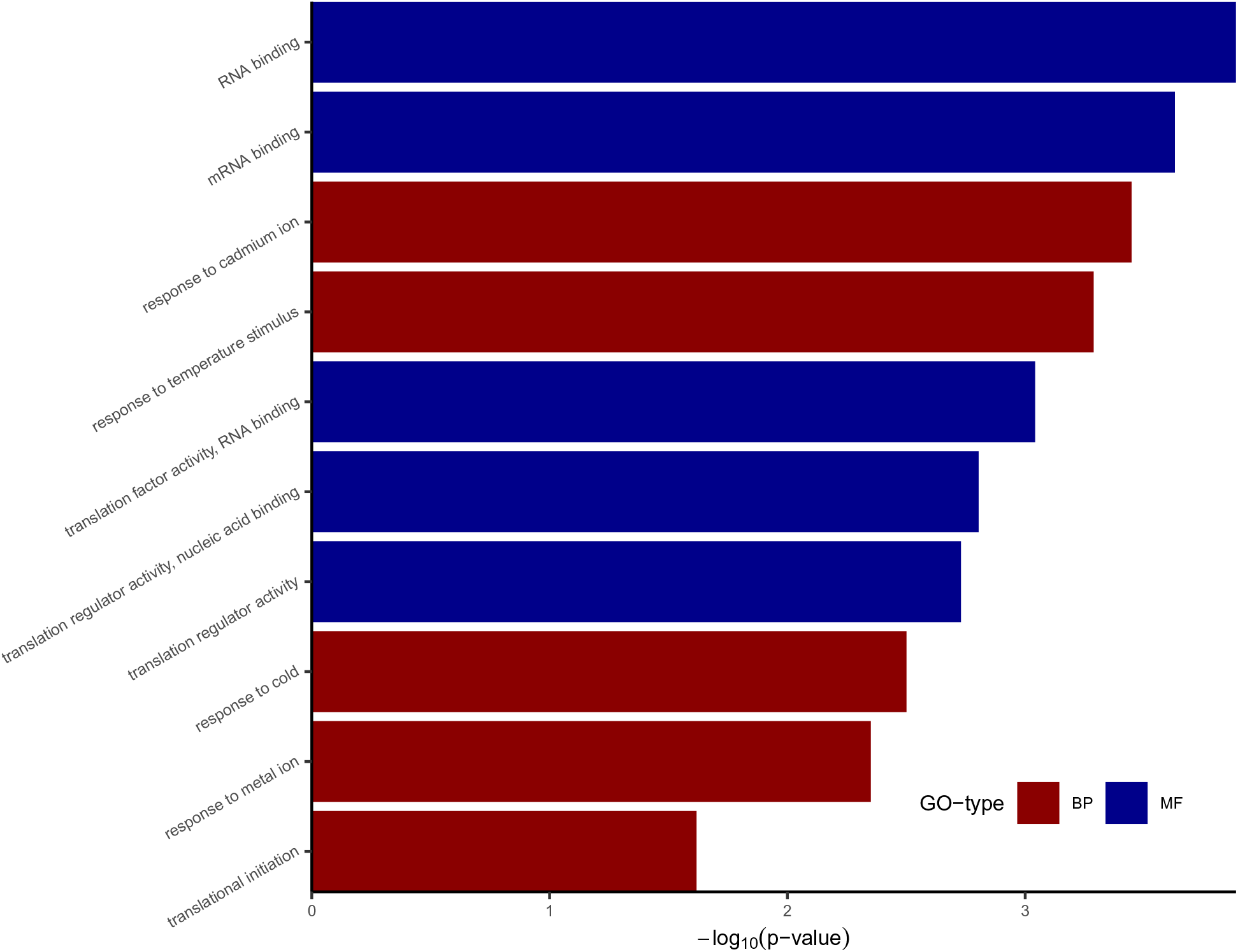
GO-enrichment of predicted signaling peptides differentially expressed between WT and *top1top2*. The labels correspond to the GO-term; red labels show biological processes while blue shows molecular function. The x-axis indicates the number of genes in the category, and the y-axis shows the -log10(p-value) of the statistical test for overrepresentation.

### Predicted TOP-regulated bioactive peptides cause unique patterns of perturbation of ETI phenotypes

Next we investigated predicted TOPs substrates from the 60 predicted bioactive peptides. Eight TOP-regulated bioactive peptides (one TOP product and seven substrates) containing at least one predicted cleavage site, were identified (**Supplemental Table S5**). The peptides mapped to six proteins: APX1, DXR, ELF5A-2/FBR12, AT5G08670, EF1A and PSBR. We next designed a peptide bioactivity validation set by adding peptides with the high differential accumulation in the peptidome and with cleavage products with significant bioactivity (**Supplemental Table S6**). The screened peptides (one TOP product and seven TOP substrates) mapped to seven proteins: APX1, DXR, FBR12, AT5G08670, PSBR, ATPD and KIN2. Two peptides mapped to APX1 while all other proteins contained only one putative bioactive TOP substrate. We also included an extended version of the shortest peptide mapping to DXR to a 12 AA peptide with predicted bioactivity. As controls, we added two peptides mapping to RPS10C and GAPA2 with no predicted bioactivity. Finally, we added two predicted bioactive peptides from proteins EIF4A1 and GSTF8 (which do not contain TOP cleavage sites) as controls for assesing TOPs regulatory role.

We examined the potential of our predicted peptide products as phytocytokines, peptides with immunoregulatory functions (Hou et al., 2021). We screened all TOP-regulated bioactive peptides and controls using a flood innoculation assay to quantify their effect on the plant ETI phenotype. For this, peptides were synthesized, dissolved in plant growth media to a concentration of 100 nM, and used to treat Arabidopsis seeds as described in the *Peptide treatments* methods section (Zipfel et al., 2006; Chinchilla et al., 2007). For the pathogen inoculation assays, two-week-old Arabidopsis seedlings with six to eight rosette leaves were grown on a solid growth medium inoculated with *Pst*AvrRpt2. Control plants were inoculated with *Pst*AvrRpt2 without peptide added to the growth medium. Colony-forming units (CFU) were measured at 0 days post inoculation (dpi) and 3 dpi and normalized for plant weight to assess the impact of peptide treatments on plant ETI response to *Pst*AvrRpt2 (**Supplemental Table S7**).

At 3 dpi, most peptide treatments significantly increased WT susceptibility to *Pst*AvrRpt2 infection (**Figure 9, Supplemental Figure S12**). For five of the tested peptides, IKTDKPFGIN (PSBR), SVVKLEAPQLAQ (ATPD), PTSTGAAKAVALV (GAPA2), ASIKVHGVPMSTA (GSTF8) and AGSAPEGTQFDARQF (EIF4A1) the treatment had no differentiating effect between *top1top2* and WT. Peptide ASIKVHGVPMSTA from the GSTF8 redox enzyme, which we predicted as bioactive but not a TOPs substrate or product, significantly increased plant susceptibility to *Pst*AvrRpt2 in WT and *top1top2*, supporting its TOP-independent role in ETI. A minor decrease in *top1top2* susceptibility compared to WT, at 3 dpi, was recorded for the peptide AGPRAGGEFGGEKGGAPA mapping to 40S ribosomal protein RPS10C.

**Figure 9:**
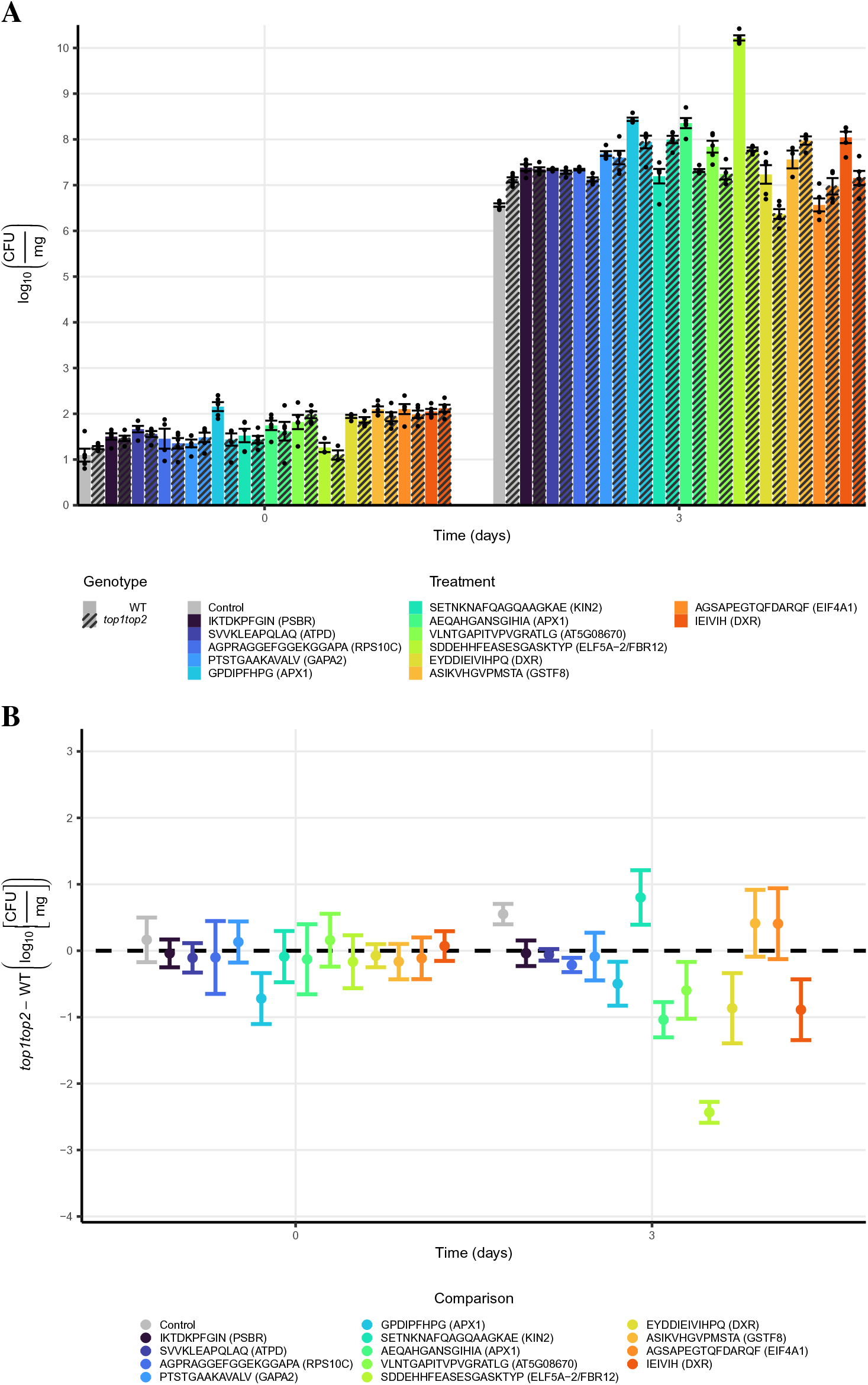
Peptide treatment causes perturbations in bacterial CFU during ETI. Colors indicate the peptide treatment: *control* or *peptide sequence* (with the corresponding protein in parenthesis) for plants grown without/with peptide treatment. The x-axis shows the measurement time. The y-axis in **(A)** shows the mean value of five replicates in log_10_(CFU/mg); error bars correspond to one standard error of the mean, and each black dot indicates one measurement. (**B**) shows the distribution of the test statistic between *top1top2* and WT. The y-axis shows the difference in mean between the genotypes with a 95 % confidence interval inferred from an equal variance t-test. The dashed line marks the null hypothesis of the test statistic. A confidence interval without overlap with the null hypothesis is significant at the 0.05 significance level.

Several peptides significantly affected bacterial growth at 3 dpi in *top1top2* relative to WT. The KIN2-derived peptide SETNKNAFQAGQAAGKAE increased susceptibility to *Pst*AvrRpt2 in *top1top2*. The KIN2 (a stress-induced gene with a cell-to-cell mobile mRNA) peptide was independently predicted as bioactive by PeptideLocator (Mooney et al., 2013). SETNKNAFQAGQAA, another putative TOPs cleavage product of KIN2, was also identified as bioactive by PeptideLocator. Plant treatments with SETNKNAFQAGQAAGKAE caused a significant susceptibility increase only in the *top1top2* line. Conversely, treatments with peptides VLNTGAPITVPVGRATLG from the ATP synthase β-subunit, AEQAHGANSGIHIA and GPDIPFHPG from APX1, IEIVIH/EYDDIEIVIHPQ from DXR, and SDDEHHFEASESGASKTYP from FBR12, resulted in significantly lower CFU counts in *top1top2* mutant. Treatments with the two APX1-derived peptides had some of the strongest effects on plants. AEQAHGANSGIHIA and GPDIPFHPG strongly increased *Pst*AvrRpt2 growth in WT but had a comparatively lower effect in the *top1top2* background. Notably, GPDIPFHPG was predicted to be bioactive, a TOPs substrate, and accumulated in *top1top2* at all timepoints. TOPs cleaved peptide products from GPDIPFHPG were also predicted as bioactive, which may account for the strong perturbation of both genotypes (third largest in WT). Notably, GPDIPFHPG and one of its TOPs-cleaved products were independently predicted as bioactive by PeptideRanker (Mooney et al., 2012). AEQAHGANSGIHIA, predicted to be bioactive and a TPP (accumulated in WT but not *top1top2* at 30 mpi), caused a severe infection in WT but only a minor increase in *top1top2* bacterial growth. Interestingly, for both APX1 peptide treatments *top1top2* rescues the WT ETI phenotype (as evidenced by decreased CFU values), albeit with distinct strengths. Together, these results argue for a substantial regulatory role of TOPs in ETI redox response through APX1 proteolysis.

The most substantial effect on *Pst*AvrRpt2 growth came from SDDEHHFEASESGASKTYP, designated as a TOP substrate and derived from the Arabidopsis eukaryotic translation initiation factor 5A (ELF5A/FBR12). Treatment with SDDEHHFEASESGASKTYP caused extreme susceptibility in WT (CFU increase of ∼ four orders of magnitude over control), which was rescued to WT levels in *top1top2*. Another predicted bioactive peptide, IEIVIH, mapped to DXR from the methylerythritol 4-phosphate pathway for isoprenoid biosynthesis. Since IEIVIH was predicted to be a TOP substrate due to its over-accumulation in *top1top2*, we also screened a longer TPS peptide with a higher bioactivity score but not detected in our peptidome (EYDDIEIVIHPQ). Interestingly, both IEIVIH and EYDDIEIVIHPQ increased WT susceptibility, with IEIVIH having a comparatively more substantial effect on CFU growth. On the other hand, *Pst*AvrRpt2-inoculated *top1top2* was insensitive to IEIVIH or, in the case of EYDDIEIVIHPQ, had increased resistance compared to WT (**Figure 9, Supplemental Figure S12**). Finally, we tested VLNTGAPITVPVGRATLG mapping to mitochondrial ATP synthase β-subunit and found that it caused increased susceptibility in WT but reduced bacterial growth in *top1top2*. This peptide accumulated in *top1top2* at 30 mpi, correlating well with the loss of proteolytic activity in the double mutant. Further, its accumulation coincided with the peak of ATP deficiency in *top1top2* at 30 mpi.

All screened peptides, except AGSAPEGTQFDARQF, significantly increased susceptibility in WT; in *top1top2* the effects were mixed, with several peptides rescuing the bacterial growth phenotype to CFU levels lower than control (**Supplemental Figure S12**). Three of the screened peptides that mapped to APX1, AT5G08670, and DXR, significantly increased susceptibility in WT without any significant effect on *top1top2*, suggesting that TOP cleavage products negatively regulate ETI.

## Discussion

Two main concepts dominate our understanding of proteases and peptidases: (1) proteolysis as an irreversible degradative mechanism of proteins required to maintain physiological homeostasis and (2) proteolysis as a targeted process with regulatory impacts on plant physiology. Unbiased studies characterizing peptidomes in complex biological samples and mutant backgrounds are necessary yet still scarce in plant systems. We describe a quantitative mass spectrometry-based peptidomics and computational approach to characterize TOPs’ proteolytic activity and its impact on the immune response. We provide temporal snapshots of the Arabidopsis peptidome following infection with an avirulent pathogen. This strategy facilitated the identification and validation of TOPs substrates, analysis of functional relevance of predicted TOPs cleavage products, and generation of an *in-silico* model for TOPs proteolytic activity.

We quantified the peptidomes in WT and *top1top2* mutant and performed high-throughput data mining to identify the cleavage patterns of TOPs and select potential bioactive peptides associated with TOPs activity. We measured the dynamics of the differential peptidome at 0 mpi and two critical time points during the immune response to *Pst*AvrRpt2. The differential peptidome at 0 mpi indicated that there are peptides associated with transcription regulatory processes, while the peak of the differential peptidome was at 30 mpi, showing significant changes in the TOP-related peptidome early in the ETI response. Functional analysis revealed that differential peptides at 0 mpi were enriched in photosynthetic processes. In comparison, metabolic processes, including ATP synthesis, glycolysis, and translation regulation (ribosome biogenesis and plastid translation), were enriched at 30 mpi. We discriminated TOP-cleaved peptides from the total complement of peptides processed during ETI through a multi-stage process, whereby we used *in vitro* validated TOP substrates from our previous screens (Al-Mohanna et al., 2021; Iannetta et al., 2021), as well as discovered new TOP substrates and analyzed the cleaved sequence patterns on all confirmed substrates. We used these insights to develop a support vector machine computational method that learned to differentiate cleavage patterns of validated substrates from the background peptidome and validated negative examples.

The study of peptidase regulatory functions is challenging due to overlap in substrate specificity (necessary redundancy for critical degradome outcomes) as well as broad substrate recognition. Despite the broad substrate selection characteristic of metallopeptidases, patterns of specificity have been described for prokaryotic and metazoan peptidases (Oliveira et al., 2001; Berti et al., 2009; Lim et al., 2007; Rawlings et al., 2018). Previous studies have shown that Arabidopsis and *Homo sapiens* TOPs and other related metallopeptidases do not have strict cleavage specificities (Tavormina et al., 2015; Polge et al., 2009; Al-Mohanna et al., 2021). As discussed in a recent review (Ferro et al., 2020), peptides generated as part of human THOP1 proteolytic pathways have many functions in immunity, signaling, and control of biological processes, including transcription regulation and metabolic regulation. Our analysis identified a distinct cleavage pattern for TOPs, LX↓XP (where L is large, P is proline, and X is any other property), including low frequency, large, hydrophobic amino acids alongside Pro. Our computational model predicts that 35 sites carrying this motif in the peptidome are cleaved (making up 8.6% of the TOPs predicted cleavage sites), while the remaining 57 were not. While the percentage of motif-carrying cleaved sites is very high, other determinants, such as PTMs or the composition of the remaining sequence, could account for the difference. Predicted cleavage sites had ∼12 % less negative amino acids (AA), ∼10 % smaller AA, ∼10 % more positive AA, and 5 % less hydrophobic AA compared to sites without a cleavage prediction. Among the 35 predicted motif-carrying cleavage sites, five are in peptides that exhibit differential accumulation in the mutant, including a) MDSDFG↓IPR from protein PFP-ALPHA2 (Q9C9K3), a regulatory subunit of pyrophosphate-fructose 6-phosphate 1-phosphotransferase (validated *in vitro*), b) (SKY)G↓SPRIVNDG (parenthesis indicating extension from protein sequence) from protein CPN60B1 (P21240), the chaperonin 60 subunit beta 1, c) VDSVFQ↓APMGTGTHH, from RCA (P10896), a protein with a role in activating RuBisCO, d) VGSFE↓SPKLSSDTK, from PUB16 (Q9LZW3), an E3 ubiquitin ligase, and e) GSSFL↓DPK, from protein PSBO1 (P23321), the oxygen-evolving enhancer protein 1-1 (validated *in vitro*). Less restricted cleavage patterns also correlate well with differential peptide accumulation. There were 60 predicted cleavage sites with motif LX↓XX (1280 not cleaved), with 25% differentially accumulating in *top1top2* and 176 predicted cleaved sites carrying the motif XX↓XP (2968 not cleaved), with 55% of these differentially accumulating in *top1top2*. These observations reinforce that Pro in the P2’ position is a strong determinant for TOP proteolytic activity and has a putative proteolytic regulatory function. GO analysis of predicted substrates further supports the idea that TOPs regulatory effect is related to redox processes and translation regulation (**Figure 3**). Thus computational xmodeling has enabled our identification of a specific TOP cleavage motif by extracting the support vectors. Applying the model, we could discriminate the TOPs-cleaved peptides from the proteolytic activity of complementary peptidases. Despite their low occurrence, large amino acids gave the model strong discriminative power, similar to the motifs for other M3 metallopeptidases in the MEROPS database. The preference for Pro at P2’ and Phe at P2 mirrors the [G/**F**k/spg/**P**fr↓**F**sr/rp/Q/] cleavage pattern reported in M3 metallopeptidases (Rawlings, 2016).

Another essential point to emerge from this study is the prediction of bioactive peptides from a large-scale peptidomics screen using a newly derived bioinformatics method. Various machine learning techniques have been used to predict peptide bioactivity (Nardo et al., 2018; Mooney et al., 2013, 2012). In plants, limited data is available to learn bioactive peptide characteristics, making it challenging to predict patterns of bioactive peptides. Here we targeted *de novo* identification of bioactive TOPs substrate peptides. A Markov chain model discriminated peptides with distinct amino acid sequences (least frequent amino acid chains) in the measured peptidome. Of note, most bioactive peptides reported in a recent review (Chen et al., 2020) score high with our model, with half of them scoring significantly above average. Intersecting the least frequent chain peptides with TOPs substrates accumulating in *top1top2* predicted potential TOPs-generated bioactive peptides. These predicted peptides mapped to regulatory enzymes from redox and ATP metabolism, such as APX1 (L-ascorbate peroxidase 1), DXR (1-deoxy-D-xylulose 5-phosphate reductoisomerase), FBR12 (eukaryotic translation initiation factor 5A-2), and several ATP synthase subunits.

We validated the ETI-related bioactivity of several peptides mapping to redox modulatory enzymes using *Pst*AvrRpt2 infection assays (**Supplemental Table S7**). Significant defects in ETI occurred when plants were grown in the presence of selected peptides (**Figure 9**). In particular, we measured a significant increase in bacterial growth for WT plants grown in the presence of APX1 peptides, indicating that APX1 turnover can modulate the ETI response. GPDIPFHPG, a TOPs substrate, consistently accumulated in the *top1top2* during the early stages of ETI. Since the peptide-treated WT had increased susceptibility, an impaired TOP-dependent proteolytic cascade could likely explain the slow ETI response of *top1top2* (Westlake et al., 2015). Accumulation of APX1 cleavage product AEQAHGANSGIHIA may also inhibit ROS-mediated signaling during ETI, increasing plant susceptibility. Notably, the most considerable effect of a peptide treatment was measured for a peptide from the FBR12 translation initiation elongation factor, providing a link between TOPs proteolytic activity and translational control, supported by the overrepresentation of the related GO category (**Figure 3B**). The plant ELF5A-2 is critical in plant growth (Feng et al., 2007) and for the PCD triggered by bacterial pathogens (Hopkins et al., 2008). Our discovery of a putative bioactive peptide mapping to FBR12 suggests a new TOPs-related PCD regulatory pathway in ETI. Two putative bioactive peptides mapped to the NAPH consumer enzyme DXR, part of the MEP pathway that provides the basic five-carbon units for isoprenoid biosynthesis (Carretero-Paulet et al., 2002). This suggests a link between TOPs proteolytic activity and regulation of isoprenoid biosynthesis, which regulates biosynthetic pathways of GA and ABA (Xing et al., 2010). Finally, a validated peptide mapping to mitochondrial ATP synthase β-subunit, a catalytic subunit with roles in plant stress (Zancani et al., 2020), directly links TOPs proteolytic activity and the ATP burst dynamics observed in ETI.

### TOP role in the regulation of ROS signaling and metabolic homeostasis

An important finding from our study of peptidome dynamics regulated by TOPs is related to the functional role of peptidases in ETI. ROS accumulation elicited by ETI is biphasic with a low amplitude and transient first phase, followed by a sustained phase of much higher magnitude (Lamb and Dixon, 1997). The degradation of peptides could require higher ROS levels produced in the second oxidative burst phase. We propose that TOPs maintain proteostasis within specific processes such as photosynthesis and ATP synthesis as ETI progresses. We discovered APX1 peptides as putative TOP targets and bioactive peptides. Hence, APX1 turnover—and consequently accumulation of APX1 bioactive peptides—may be essential in monitoring ROS scavenging and production.

In mammalian systems, the hormonal peptide atrial natriuretic peptide (ANP) activates NOX2, causing ROS accumulation (Fürst et al., 2005). Since the well-studied NADPH/respiratory burst oxidase proteins (RBHOs) are homologs of NOX2, a potential mechanism emerges whereby TOP-dependent proteolytic events influence ETI by modulating ROS signaling and PCD pathways. We found candidate cleavage sites in the cytosolic NADP-dependent isocitrate dehydrogenase (cICDH) involved in redox and metabolic control (Mhamdi et al., 2010). Further, we found several peptides from GAPDH subunits known to function in ROS regulation and plant resistance (Henry et al., 2015) and peptides from pyrophosphate-fructose 6-phosphate 1-phosphotransferase subunit alpha 2 (PFP-ALPHA2) acting in glycolysis (Lim et al., 2009). These bioactive peptides could help regulate the carbohydrate fluxes between ATP production through glycolysis and NADP production via the pentose 5-phosphate pathway.

### ATP and NADP perturbation; the cause of the delayed ROS burst in top1top2?

Cellular damage causes the release of ATP into the extracellular matrix, where it is recognized as a damage-associated molecular pattern by P2 receptor kinase 1 (P2K1) on the plasma membrane (Choi et al., 2014; Tanaka et al., 2014). This event leads to increased production of secondary messengers such as cytosolic Ca^2+^, nitric oxide, and ROS, which mediate plant defense responses (Tripathi et al., 2018). The chloroplastic NADP(H) pool is vital for stable ATP production from the photosynthetic machinery (Hashida and Kawai-Yamada, 2019). During stress, perturbations of the NADP(H) pool can be caused by changes in the Calvin-Benson cycle, leading to ROS accumulation in the chloroplast. The NADP(H) pool was strongly perturbed in *top1top2* at the onset of ETI compared to WT (**Figure 4**). Unlike WT, *top1top2* accumulated NADP(H) during the first 30 mpi of ETI, only to stabilize at 180 mpi. Likewise, WT accumulated a large pool of ATP during this same period. A logical explanation would be that in WT, the NADP(H) pool is converted to NAD, which is used for catabolic energy production (Hashida and Kawai-Yamada, 2019) and, consequently, ATP accumulation. In *top1top2*, this rapid reshuffling of metabolic resources is perturbed and possibly delayed, which would be in accordance with its delayed ROS burst during ETI (Al-Mohanna et al., 2021). Indeed, we found a substantial change in peptide accumulations at 30 mpi between the two genotypes (**Figure 2A**), and many of the proteins these peptides belonged to are involved in ATP synthesis (**Figure 3B**). Further, *top1top2* had a decreased accumulation of NADP(H) at 0 mpi, and peptides from proteins associated with the electron transport chain unique to 0 mpi were enriched. The electron transport chain is a key driver of NADP(H) turnover in the chloroplast (Hashida and Kawai-Yamada, 2019). Thus, TOPs could play a principal role in controlling early metabolic fluxes during ETI; in *top1top2* background, deficient ROS accumulation and, consequently, ROS signaling could help explain the perturbation of its redoxome during the later stages of ETI (McConnell et al., 2019).

In conclusion, the data presented here provide a comprehensive view of peptide processing events associated with plant immune response and insights into the proteolytic activities and substrates of metallopeptidases. To our knowledge, this is the first peptidomics screen that successfully identified bioactive peptides and ETI pathways modulated by M3 oligopeptidases. Our results reveal a pattern of peptide bioactivity, arguing that TOPs are components in a complex regulatory network of peptide substrates and products generated during ETI. We show that *(1)* peptides derived from TOPs proteolytic activity increase susceptibility to a bacterial pathogen in WT while rescuing the ETI phenotype in the double mutant, *(2)* redox processes activated during ETI are likely controlled via phytocytokines produced by controlled proteolysis, and *(3)* predictive modeling methods combined with experimental validation facilitates the discovery of novel bioactive peptides. Further work on integrating targeted and systems biology approaches is key to gaining insights into protease/peptidase networks in relevant biological models to understand their role in mediating signaling and coordination with other types of regulatory protein modifications.

## Experimental Procedures

### Plant Growth and Infection Assays for ETI peptidomics screen

Seeds of *Arabidopsis thaliana* ecotype Columbia (Col-0) and *top1top2* were sterilized by standardized methods as described in (Lindsey III et al., 2017) and grown on MS media for 10 days, then transferred to individual jiffy pellets under controlled conditions with a 12h light (12:00 pm to 12:00 am; 100 µmol m^-2^ s^-1^ photon flux density), 12 h dark period and relative humidity of 60% to 65%. Day and night temperatures were set to 23°C and 21°C, respectively. Experiments were performed with 4 to 5-week-old, uniform appearance naïve plants. *Pseudomonas syringae* pv. *tomato* pathovar DC3000 (*Pst*) carrying avrRpt2 were cultivated at 28°C in King’s B medium (Sigma Aldrich) containing Rifampicin and Kanamycin (Whalen et al., 1991). Overnight log-phase cultures were diluted to final optical densities of 600 nm (OD_600)_ for leaf inoculations of WT and *top1top2* plants. To activate ETI, two to three mature leaves at similar developmental stages were infiltrated with *Pst*AvrRpt2 suspensions in 10 mM MgCl_2_ buffer at 5 × 10^5^ CFU ml^-1^; control plants were infiltrated with 10 mM MgCl_2_. Infiltrated leaves were harvested at the required time points for peptidome analysis.

### Peptide Extraction

Three biological replicates were used for each genotype (*i.e.,* WT and *top1top2* mutant) and infection timepoint. The preparation of peptidome samples from local tissue followed the method described in (Iannetta et al., 2021). Briefly, rosette leaf tissue was ground under liquid N_2_, and peptides were extracted from plant material in two rounds using 10% trichloroacetic acid (TCA) in acetone. The isolation of peptides from small molecules in this extract was performed using strong cation exchange solid-phase extraction (SPE), and peptides were desalted using reversed-phase SPE. Peptide concentrations were estimated using the Pierce Quantitative Colorimetric Peptide Assay (Thermo Fisher Scientific) according to the manufacturer’s protocols. Based on these results, peptide concentrations in each experiment were normalized across replicates before LC-MS/MS analysis.

### LC-MS/MS Analysis

Samples were analyzed using an Acquity UPLC M-Class System (Waters) coupled to a Q Exactive HF-X mass spectrometer (Thermo Fisher Scientific). Mobile phase A consisted of water with 0.1% formic acid (Thermo Fisher Scientific), and mobile phase B was acetonitrile with 0.1% formic acid. Injections (1 μL) were made to a Symmetry C18 trap column (100 Å, 5μm, 180μm x 20 mm; Waters) with a flow rate of 5 μL/min for 3 min using 99% A and 1% B. Peptides were then separated on an HSS T3 C18 column (100 Å, 1.8μm, 75μm x 250 mm; Waters) using a linear gradient of increasing mobile phase B at a flow rate of 300 nL/min. Mobile phase B increased from 5% to 40% in 90 min before ramping to 85% in 5 min, where it was held for 10 min before returning to 5% in 2 min and re-equilibrating for 13 min. The mass spectrometer was operated in positive polarity, and the Nanospray Flex source had spray voltage floating at 2.1 kV, the capillary temperature at 320 °C, and the funnel RF level at 40. MS survey scans were collected with a scan range of 350 – 2000 *m/z* at a resolving power of 120,000 and an AGC target of 3 x 10^6^ with a maximum injection time of 50 ms. A top 20 data-dependent acquisition was used where HCD fragmentation of precursor ions having +2 to +7 charge state was performed using a normalized collision energy setting of 28. MS/MS scans were performed at a resolving power of 30,000 and an AGC target of 1 x 10^5^ with a maximum injection time of 100 ms. Dynamic exclusion for precursor *m/z* was set to a 10 s window.

### Database Searching and Label-Free Quantification

Acquired spectral files (*.raw) were imported into Progenesis QI for proteomics (Waters, version 2.0). Peak picking sensitivity was set to the maximum of five, and a reference spectrum was automatically assigned. Total ion chromatograms (TICs) were then aligned to minimize run-to-run differences in peak retention time. Each sample received a unique factor to normalize all peak abundance values resulting from systematic experimental variation. Alignment was validated (>80% score), and a combined peak list (*.mgf) was exported out of Progenesis for peptide sequence determination by Mascot (Matrix Science, version 2.5.1; Boston, MA). Database searching was performed against the *Arabidopsis thaliana* UniProt database (https://www.uniprot.org/proteomes/UP000006548, 39,345 canonical entries, accessed 03/2021) with sequences for common laboratory contaminants (https://www.thegpm.org/cRAP/, 116 entries, accessed 03/2021) appended. Target-decoy searches of MS/MS data used “None” as the enzyme specificity, peptide/fragment mass tolerances of 15 ppm/0.02 Da, and variable modifications of N-terminus acetylation, C-terminus amidation, and methionine oxidation. Significant peptide identifications above the identity or homology threshold were adjusted to less than 1% peptide FDR using the embedded Percolator algorithm (Käll et al., 2007). Mascot results (*.xml) were imported to Progenesis for peak matching. Identifications with a Mascot score less than 13 were removed from consideration in Progenesis before exporting both “Peptide Measurements” and “Protein Measurements” from the “Review Proteins” stage.

### LC-MS/MS data analysis - peptidomics

Data were parsed using custom scripts written in R for pre-processing and statistical analysis (https://github.com/hickslab/QuantifyR). The “Peptide Measurements” data contain peak features with distinct precursor mass and retention time coordinates matched with a peptide sequence identification from the database search results. Some features were duplicated and matched with peptides having identical sequences, modifications, and scores but alternate protein accessions. These groups were reduced to satisfy the principle of parsimony and represented by the protein accession with the highest number of unique peptides found in the “Protein Measurements” data for this experiment else, the protein with the largest confidence score was assigned by Progenesis. Some features were also duplicated with differing peptide identifications and were reduced to just the peptide with the highest Mascot ion score. An identifier was created by joining the protein accession of each peptide to the identified peptide sequence. Each dataset was reduced to unique identifiers by summing the abundance of all contributing peak features (*i.e.,* different peptide charge states and combinations of variable modifications). Identifiers were represented by the peptide with the highest Mascot score in each group. Identifiers were removed if there was not at least one condition with > 50% nonzero values across the abundance columns.

### Differential peptidomics analysis

To compare peptide abundance between treatments, we used the linear model analysis from *limma* (Law et al., 2014; Ritchie et al., 2015) with the mean variance trend correction described in the next section. We first filtered out peptides with more than 15 missing values; then, all samples were mean stabilized using the size factor normalization described in (Anders and Huber, 2010) and were log_2_ transformed (**Supplemental Figure S13**). After data normalization, multiple imputations were performed using the imputation model described below in *Data imputation*. We ran 1500 imputations and used a p-value cut-off of 0.05 and |logFC|≥2, while the p-value cut-off for the binomial test with null hypothesis p>0.5 was set to 0.05. After analysis, sets of differentially abundant peptides were compared and visualized with UpSetR (Conway et al., 2017).

### Mean-variance trend correction

We model the standard deviation of peptides as a *gamma* function of the mean:

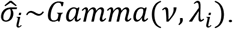

*Gamma* regression (Nelder and Wedderburn, 1972) was used to model dependency between peptide abundance variation and its mean:

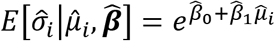

where *μ̂_i_* and *σ̂_i_*, are the sample mean and standard deviation for the i: the peptide.

To remove the mean-variance trend in the data, we estimated precision weights using the *gamma* regression. The corresponding precision weight, *w_ijr_*, of the data-point *x_ijr_* (i:th peptide in the j:th condition for the r:th replicate), was calculated as the squared inverse of the expected value of the standard deviation given the data-point and the estimated regression coefficients:

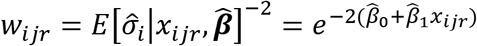

The parameters for the regression model were estimated using the *glm* function in R (RC team 2013, version 4.0.5) with the flag *family=Gamma(‘log’)*.

### Data imputation

We used a normal distribution as missing data imputation model:

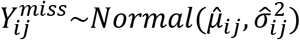

To estimate the mean of the normal distribution for the i:th peptide in the j:th condition, *μ̂_ij_*, we used the sample mean of the non-missing data points:

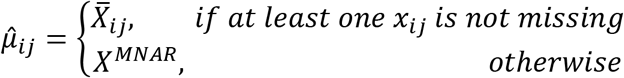

If all data points were missing, we assume that the data was missing not at random (MNAR) due to the concentration of the peptide being below the level of detection. For these cases, we used Tukey’s lower fence as mean estimate:

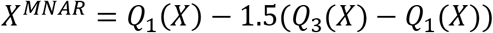

To estimate the standard deviation, *σ̂_ij_*, we used one gamma regression per condition to find the expected standard deviation value given the peptide mean and the regression coefficients for the condition.

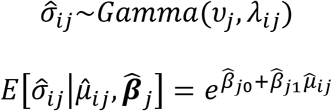

The regression parameters were estimated as for the mean-variance trend correction.

### GO analysis

The UniProt accessions of the proteins mapping DAPs were converted to Araport IDs using UniProt’s ID mapper. Gene ontology (GO) annotations were assigned from UniProt. GO term enrichment was performed using the ThaleMine overrepresentation testing (Fisher’s Exact with false discovery rate correction).

### SVM-model proteolysis prediction

We used a support vector machine (SVM) with a Laplace kernel and a least-squares estimator (duVerle and Mamitsuka, 2012) to find model parameters. From motif analysis, we found that the six positions symmetrically arranged around the cleavage site provide most of the predictive model (**Figure 5**). As before (see section *Predictive model of TOP peptidases proteolytic activity*), all peptides were extended by adding an “e” at the N-terminus, symbolizing that the positions were empty. Training data was generated by sliding the six-position window over all validated peptides and the negative examples. In total, this generated 528 training examples, 49 positive/cleaved and 479 negative/not cleaved. Due to the large discrepancy between positive and negative examples, positive examples were weighted. To determine the weights, we performed 10-fold cross-validation five times. We found that weighting the positive examples four times in the training procedure maintained a low false-positive rate and high true negative rate (**Supplemental Figure S14**) while generating a higher true-positive rate. Model training was performed in R (RC team 2013, version 4.0.5) using the *lssvm* function from the kernlab package (Karatzoglou et al., 2004) with the flags kernel=’laplacedot’, centered = F, kpar = list(sigma =0.1288), and tol = 10^-150^ and prediction was done using R’s predict function.

### Motif Analysis

In preparation for sequence logo visualization, data were parsed using custom scripts written in R. Peptides were filtered for those significantly increasing in WT (*p*-value < 0.05, log_2_-transformed fold change ≤ -1). Peptide termini were extended with four amino acids from corresponding protein sequences or until the protein terminus was reached. Each elongated peptide was truncated into two smaller peptides containing the extended amino acids and either the first or the last four amino acids of the original peptide sequence. These truncated peptides were filtered for those with eight residues to satisfy the input condition requiring peptides of the same length. For motif analysis, sequence logo visualizations were performed using pLOGO; positions with significant residue presence are depicted as amino acid letters sized above the red line (O’Shea et al., 2013).

### Cleavage Motif Analysis

To determine the overrepresentation of amino acids/properties (section *Characterization of TOPs cleavage patterns*), we used the pLogo software (O’Shea et al., 2013) without the option *remove duplicate sequences*. For the background, the windows without a predicted cleavage site from the SVM model were used. The foreground used is described by the context in the Results section. Fasta files inputed to pLogo was produced using the in the supplemental R code archive.

### Bioactive Peptide Prediction Model

Let 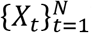 be a sequence of random variables that describe amino acid patterns in the primary structure of proteins (Onicescu, 1977). We define a twenty-state Markov Chain (MC) over all sequential pairs of amino acids observed in a peptidome dataset. That is, if the dataset had two proteins with amino acid sequences ALLA and LLAA, then all transitions would be AL, LL, LA, LL, LA, and AA, where the first three are from the first protein and the fourth to the sixth from the second. We computed all MC parameters over the ETI peptidome dataset and used the Araport11 proteome as a reference (Cheng et al., 2017).

Let *p_ij_* be the transition probabilities of the MC going from the i:th to j:th amino acid. We used the maximum likelihood estimator to infer the transition probabilities:

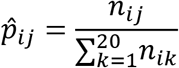

where *n_ij_* is the observed number of transitions from the i:th to j:th amino acid. We define the uniqueness of an amino acid sequence as:

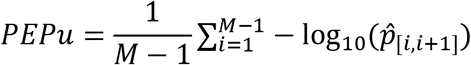

where M is the number of amino acids in the peptide and *p̂*_[*i,i*+1]_ is the probability of the i:th transition. The uniqueness measure of all peptides in the dataset was then standardized:

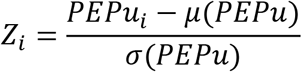

(where *μ*(.) and *σ*(.) are the estimated mean and variance) and a right-tailed Z-test was performed to generate a p-value for each peptide:

### In Vitro Enzyme Assay

Heterologously expressed and purified TOP enzymes, and synthetic peptides were produced as described in our previous work (Iannetta et al., 2021). Synthesized peptides were solubilized in 500 µL of 100 mM NaCl in 50 mM Tris, pH 7.5. To initiate the enzyme assay, either TOP1, ChlTOP1, or TOP2 was added at a peptide:TOP ratio of 10:1. The reaction mixture was incubated at 23 °C for 30 min. Reaction mixtures were desalted using reversed-phase SPE.

### LC-MS Analysis – in vitro enzyme assays

*In vitro* enzyme assay samples were analyzed using an LC-MS/MS platform, as previously described (Kirkpatrick et al., 2017), with the following specifications: 0.1% formic acid in all mobile phases and a trapping mobile phase composition of 1% acetonitrile/0.1% formic acid. The MS was operated in positive-ion, high-sensitivity mode with the MS survey spectrum using a mass range of *m/z* 350−1600 in 250 ms and information-dependent acquisition of MS/MS data using an 8 s dynamic exclusion window. The first 20 features above an intensity threshold of 150 counts and having a charge state of +2 to +5 were fragmented using rolling collision energy (CE; ±5%).

### Quantitative ATP Luciferase Assay

Four biological replicates of frozen rosette leaf tissue (∼0.05 g) were pulverized via three, 5 min rounds using a TissueLyser II (Qiagen, Germantown, MD) cell disrupter at 30 Hz before homogenization in 250 μL of water. The homogenate was vortexed and incubated at 100 °C for 30 min before centrifugation at 10,000 *g* for 15 min at 4 °C. The supernatant was collected, protein concentrations were estimated using the CB-X assay (G-Biosciences, St. Louis, MO) according to the manufacturer’s protocol, and these concentrations were used to normalize replicates following luminescence analysis. The ATP determination kit (Invitrogen, Waltham, MA) was used to quantify cellular ATP concentrations according to the manufacturer’s protocol. Briefly, 20 μL of sample and 180 μL of 1 mM DTT, 0.5 mM luciferin, and 1.25 μg/mL firefly luciferin in 1x reaction buffer were added to a 96-well plate and incubated in the dark for 5 min at RT. Luminescence was measured and compared to an ATP standard calibration curve to convert luminescence to ATP concentration. Each biological replicate was analyzed in technical triplicates.

### Quantitative NADP^+^/NADPH Enzyme Cycling Assay

The NADP/NADPH Quantitation Kit (Sigma-Aldrich, Burlington, MA) was used to quantify cellular NADP^+^/NADPH concentrations according to the manufacturer’s protocol. Briefly, five biological replicates of frozen rosette leaf tissue (∼0.05 g) were pulverized via three 5 min rounds using a TissueLyser II (Qiagen) cell disrupter at 30 Hz before homogenization in 500 μL of the provided extraction buffer. The homogenate was vortexed and incubated at −20 °C for 10 min before centrifugation at 10,000 *g* for 15 min at 4 °C. The supernatant was collected and filtered through a 10 kDa molecular weight cutoff filter by centrifugation at 3,200 *g* for 30 min at 4 °C. Prior to filtering, an aliquot was reserved for protein quantification via the CBX assay (G-Biosciences); this protein concentration was used to normalize NADP^+^/NADPH based on protein quantification following analysis. For total NADP^+^/NADPH quantification, 200 μL of the filtrate was incubated for 30 min at 60 °C before centrifugation at 10,000 *g* for 5 min at 4 °C. After incubation, 10 μL of developer solution was added, and mixtures were incubated in the dark for 30 min at RT. Absorbances were measured (λ = 450 nm) and compared to an NADP^+^/NADPH standard calibration curve to convert absorbance to NADP^+^/NADPH concentration. Each biological replicate was analyzed in technical duplicates.

### Peptide treatments

Selected peptides were synthesized via Fmoc-based solid-phase peptide synthesis using a flow chemistry-based platform, which was built in-house based on the prior invention (Mijalis et al., 2017; Simon et al., 2014), and lyophilized. The molecular weights of the peptides were calculated using their sequences (https://www.bioinformatics.org/sms/prot_mw.html). Lyophilized peptide powder was measured with a digital analytical balance (Accuris instruments, W3100 series) and resuspended in a sterile buffer containing 100 mM NaCl (Sigma Life Sciences), 50 mM Tris (Fisher Scientific) at pH 7.5. For assays, the peptide solutions were further diluted to 100 nM aliquots to avoid freeze-thaw cycles and stored at −20°C before use. Arabidopsis seedlings were grown on ½ MS solid medium into 12-well plates filled halfway for flood inoculation assays. The growth medium was autoclaved and cooled to 55°C in a water bath before adding the peptide preparations.

### Flood Inoculation Assays for ETI Phenotype Screening

*Arabidopsis thaliana* seeds were sterilized with 70% (v/v) ethanol and 50% (v/v) bleach solution (Clorox, 4.5% sodium hypochlorite) followed by washing with sterilized distilled water as described in (Lindsey III et al., 2017). The seeds were transferred to square Petri dishes containing solid ½ MS medium (2.2 g/L, without vitamins) with 10 g/L sucrose (MP Biomedical, LLC, USA) and 0.3 % PhytagelTM, in the absence (control) or presence of 100 nM peptide (treatment), as described in (Ishiga et al., 2017). Then plates were transferred to 4°C for three days, then to a growth chamber with photoperiods of 12h light (100-200 µmol m-2 s-1 photon flux density) followed by 12 h dark. The growth chamber temperature was set to 23°C during the day and 21°C during the night and relative humidity of 60%-65%. The seedlings were grown for 2 weeks before inoculation. *Pst*AvrRpt2 was grown in liquid King’s medium B containing kanamycin and rifampin (50 and 25 µg/ml, respectively) and cultured in 28°C on a shaker incubator. After 8-10 hours, 1 ml bacterial culture was centrifuged, at 9600 RPM for two minutes, and the pellet was resuspended in 1 mL of 10mM MgCl_2_ buffer. The bacterial density was then adjusted to an OD600 value of 0.1 and used in serial dilution to produce a bacterial inoculum of 5×10^3^ colony-forming unit (CFU) per ml.

Flood inoculation was performed on Arabidopsis seedlings (Ishiga et al., 2011, 2017). Briefly, 0.025% Silwet L-77 (Lehle Seed, TX, USA) was added to the bacterial inoculum and mixed gently. 40 ml of bacterial inoculum was distributed in the plates containing the two-week-old seedlings and incubated for 3 min at room temperature, after which the bacterial inoculum was discarded. For 3-dpi measurements, seedlings were sterilized in 5% H_2_O_2_ (Sigma-Aldrich), followed by a wash with sterile distilled water (sdH_2_O). They were weighed and then homogenized in 300 µl sdH_2_O. Serial dilution was performed six times (for 0dpi) and ten times for (3dpi) with 10 µl sample/dilute and 90 µl sdH_2_O. The diluted cultures were plated on King Agar B (Sigma-Aldrich) containing kanamycin and rifampin (50 and 25 µg/ml, respectively) and incubated at 28°C for 2 days. The bacterial CFU was counted and normalized with the total weights of the plant tissue sample collected. Significance testing was performed using a pooled-variance t-test.

## Data Availability

The mass spectrometry peptidomics data have been deposited to the ProteomeXchange Consortium via the PRIDE partner repository (Perez-Riverol et al., 2019) and can be accessed with the dataset identifier PXD019812 and 10.6019/PXD019812.

Username

Password: IIhTfPXT

## Code availability

The peptidome normalization, imputation, and differential abundance analysis pipeline code is available at: https://github.com/PhilipBerg/pair. An archive of the R-code and the data analyzed in this paper is available for download at: https://figshare.com/s/ccda4b4596909ded3d26.

## Acknowledgments

This work was supported by the National Science Foundation (collaborative NSF-MCB 1714405/ 1714157 to S.C.P., G.V.P., and L.M.H) and a National Science Foundation Major Research Instrumentation award (CHE-1726291 to L.M.H) for the Q Exactive HF-X mass spectrometer.

## Author contributions

GVP, LH and SCP devised the project and the main conceptual idea; NN grew plants, performed all plant ETI assays, and collected the plant material for the peptidomics screen; SCP supervised the plant ETI assays and enzyme characterization assays; AAI carried out mass spectrometry experiments and analysis of resultant data and performed in vitro screening of TOPs substrates; AS performed the ATP luciferase and NADP(H) enzyme cycling assay; LMH supervised the mass spectrometry, ATP, and NADP(H) assays and analyses; AP, ZB and AW performed the peptide synthesis; PB implemented the statistical data analysis pipeline and the machine learning code; PB and GVP performed the statistical analysis of data; RS and UW performed flood inoculation assays and peptide treatments; GVP supervised the data analysis, computational modeling and peptide bioactivity validation; GVP, PB, AAI and SCP wrote the manuscript; LMH, AW, AS and RS edited and contributed to the writing. All authors discussed the results and commented on the article.

## Figure Legends

**Supplemental Figure S1:** MA-plots showing the distribution of the decision of the differential abundance analysis for the peptides in time-course analysis. The title of each facet corresponds to the decision of that comparison. Each dot corresponds to a peptide, and the x-axis shows the mean of that peptide for that comparison, while the y-axis shows the log fold-change, and a positive value indicates accumulation in *top1top2*.

**Supplemental Figure S2:** UpSetR of the significant hits in time-course analysis. It shows the size of the set(s) marked with a dot below the bars.

**Supplemental Figure S3:** Common GO terms enriched in time-course analysis. The title of the facets indicates the p-value and number of genes in the category for that comparison. The labels in the plot correspond to the different categories. The x-axis shows the number of genes in the category, and the y-axis shows the −log10(p-value) of the statistical test for overrepresentation.

**Supplemental Figure S4:** GO-enrichment of biological process unique in the time-course analysis. The x-axis shows the number of genes in the category, while the y-axis shows the -log10(p-value) of the statistical test for overrepresentation. Red indicates the GO-terms unique to 0 mpi, and blue indicates the same but for 30 mpi.

**Supplemental Figure S5: A) assay_VVISAPSKDAPM**: In vitro enzymatic assays including the synthetic peptide with the sequence VVISAPSKDAPM. (A) Extracted ion chromatograms and mass spectrum of the N-terminal product VVISAPSKD (red, m/z 915.51, +1 charge state and m/z 458.26, +2 charge state) indicate this cleavage site for TOP2. (B) Extracted ion chromatograms and mass spectra of the N-terminal product VVIS (blue, m/z 417.27, +1 charge state) indicate this cleavage site for CHLTOP1 and TOP2. All observed masses match the theoretical peptide masses within 7 ppm mass error.

**B) assay_SVVKLEAPQLAQ**: In vitro enzymatic assays including the synthetic peptide with the sequence SVVKLEAPQLAQ. (A) Extracted ion chromatograms and mass spectra of the N-terminal product SVVKLEAPQ (red, m/z 485.78, +2 charge state) indicate this cleavage site for CHLTOP1 and TOP2. (B) Extracted ion chromatograms and mass spectra of the N-terminal product SVVKLE (blue, m/z 674.41, +1 charge state) and C-terminal product APQLAQ (green, m/z 627.35, +1 charge state) indicate this cleavage site for CHLTOP1 and TOP2. (C) Extracted ion chromatograms and mass spectra of the N-terminal product SVVKL (orange, m/z 545.37, +1 charge state) indicate this cleavage site for CHLTOP1 and TOP2. All observed masses match the theoretical peptide masses within 8 ppm mass error.

**C) assay_IKTDKPFGIN**: In vitro enzymatic assays including the synthetic peptide with the sequence IKTDKPFGIN. (A) Extracted ion chromatograms and mass spectra of the N-terminal product IKTDKPF (red, m/z 848.49, +1 charge state and m/z 424.75, +2 charge state) indicate this cleavage site for CHLTOP1 and TOP2. (B) Extracted ion chromatograms and mass spectra of the C-terminal product KPFGIN (blue, m/z 675.38, +1 charge state) indicate this cleavage site for CHLTOP1 and TOP2. All observed masses match the theoretical peptide masses within 4 ppm mass error.

**D) assay_MDSDFGIPR**: In vitro enzymatic assays including the synthetic peptide with the sequence MDSDFGIPR. Extracted ion chromatograms and mass spectra of the N-terminal product MDSDFG (red, m/z 671.23, +1 charge state) indicate this cleavage site for CHLTOP1 and TOP2. All observed masses match the theoretical peptide masses within 3 ppm mass error.

**E) assay_TGDQRLLDAS**: In vitro enzymatic assays including the synthetic peptide with the sequence TGDQRLLDAS. Extracted ion chromatograms and mass spectra of the N-terminal product TGDQRLL (red, m/z 802.44, +1 charge state and m/z 401.72, +2 charge state) indicate this cleavage site for CHLTOP1 and TOP2. All observed masses match the theoretical peptide masses within 5 ppm mass error.

**F) assay_DPFGLGKPA**: In vitro enzymatic assays including the synthetic peptide with the sequence DPFGLGKPA. Extracted ion chromatograms and mass spectra of the N-terminal product DPFGLG (red, m/z 605.29, +1 charge state) indicate this cleavage site for CHLTOP1 and TOP2. All observed masses match the theoretical peptide masses within 4 ppm mass error.

**G) assay_GSSFLDPK**: In vitro enzymatic assays including the synthetic peptide with the sequence GSSFLDPK. Extracted ion chromatograms and mass spectra of the N-terminal product GSSFL (red, m/z 510.26, +1 charge state) indicate this cleavage site for CHLTOP1 and TOP2. All observed masses match the theoretical peptide masses within 2 ppm mass error.

**H) assay_VLNTGAPITVPVGRATLG**: In vitro enzymatic assays including the synthetic peptide with the sequence VLNTGAPITVPVGRATLG. (A) Extracted ion chromatograms and mass spectra of the N-terminal product VLNTGAPITVPVGRA (red, m/z 732.93, +2 charge state) indicate this cleavage site for TOP2. (B) Extracted ion chromatograms and mass spectra of the C-terminal product RATLG (blue, m/z 517.31, +1 charge state) indicate this cleavage site for CHLTOP1 and TOP2. (C) Extracted ion chromatograms and mass spectra of the N-terminal product VGRATLG (green, m/z 673.40, +1 charge state) indicate this cleavage site for CHLTOP1 and TOP2. (D) Extracted ion chromatograms and mass spectra of the N-terminal product VLNTGAPIT (orange, m/z 885.50, +1 charge state) indicate this cleavage site for CHLTOP1 and TOP2. All observed masses match the theoretical peptide masses within 7 ppm mass error.

**I) assay_AKDELAGSIQKGV**: In vitro enzymatic assays including the synthetic peptide with the sequence AKDELAGSIQKGV. Extracted ion chromatograms and mass spectrum of the N-terminal product AKDELAGSIQ (red, m/z 516.27, +2 charge state) indicate this cleavage site for CHLTOP1. All observed masses match the theoretical peptide masses within 1 ppm mass error.

**J) assay_TGGGASLELLEGKPLPG**: In vitro enzymatic assays including the synthetic peptide with the sequence TGGGASLELLEGKPLPG. (A) Extracted ion chromatograms and mass spectra of the N-terminal product TGGGASLELLE (red, m/z 532.77, +2 charge state) and C-terminal product GKPLPG (blue, m/z 568.35, +1 charge state) indicates this cleavage site for CHLTOP1. (B) Extracted ion chromatograms and mass spectrum of the C-terminal product EGKPLPG (green, m/z 697.39, +1 charge state) indicate this cleavage site for CHLTOP1. (C) Extracted ion chromatograms and mass spectrum of the N-terminal product TGGGASLE (orange, m/z 691.33, +1 charge state) indicate this cleavage site for CHLTOP1. (D) Extracted ion chromatograms and mass spectrum of the N-terminal product TGGGASL (purple, m/z 562.28, +1 charge state) indicate this cleavage site for CHLTOP1. All observed masses match the theoretical peptide masses within 8 ppm mass error.

**Supplemental Figure S6.** Sequence logo visualizations of TOP cleavage specificity using theoretical and determined TOP substrates. Positions with significant residue presence are depicted as amino acid letters sized above the red line (O’Shea et al., 2013). (A) Motif analysis of the extended termini of peptides significantly increased in the wild-type plants across the infection time points.

**Supplemental Figure S7:** Histogram of the amino acid composition and the number of observed cleavage sites of the peptides with validated cleavage sites. The y-axis shows the number of observations, and the x-axis shows the different amino acids.

**Supplemental Figure S8:** The observed number of amino acids towards the C-or N-terminus (indicated by the facet title) around the validated cleavage sites. The x-axis shows the number of positions away from the cleavage site, and the y-axis shows the different properties.

**Supplemental Figure S9:** The observed number of distinct amino acid properties towards the C- or N-terminus (as indicated by the facet title) around the validated cleavage sites. The x-axis shows the number of positions away from the cleavage site, and the y-axis shows the different properties. See section *Characterization of TOPs cleavage patterns* to describe the property categories.

**Supplemental Figure S10:** pLogo showing the overrepresentation of different amino acid properties over- or under-represented at the predicted cleavage sites. See section *Characterization of TOPs cleavage patterns* for a description of the property categories and see (O’Shea et al., 2013) for more details on interpreting the plot.

**Supplemental Figure S11:** pLogo showing the overrepresentation of different amino acids over- or under-represented at the predicted cleavage sites.

**Supplemental Figure S12:** Significant changes in bacterial growth between peptide treatment and control (no treatment) in WT and *top1top2*. Colors indicate the peptide treatment with the protein they belong to in parenthesis; the x-axis shows the measurement time; the y-axis shows the difference in mean between peptides and control with a 95 % confidence interval inferred from an equal variance t-test. The dashed line marks the null hypothesis of the test statistic. A confidence interval without overlap with the null hypothesis is significant at the 0.05 significance level.

**Supplemental Figure S13:** Median stabilization between samples from the data normalization. The y-axis shows the data values either normalized using the size factor normalization described in (Anders and Huber, 2010) in the blue boxes or the raw data after log2 transformation in the red boxes. The x-axis shows the different samples.

**Supplemental Figure S14:** Boxplots showing the results of k-fold cross-validation with different weights of the positive/cleaved examples. The titles of each facet indicate the amount of weighting given to the positive examples. The x-axis shows the evaluation metrics; FPR is the false positive rate, TNR is the true negative rate, and TPR is the true positive rate. The y-axis shows the values of the different metrics. The metrics were calculated without the weighting.

## Tables

**Supplemental Table S1:** Synthetic AtTOPs peptide substrates tested in in vitro enzyme assays. All peptides were significantly increasing in the top1top2 mutants at one or more timepoints and the order of the timepoints matches the order of the listed fold changes in column F. In columns H-I, the arrows represent identified sites of cleavage (ND: none detected) and cleaved peptide products that are bolded and underlined were uniquely detected in the enzyme-treated samples compared to the analysis of the bare synthetic peptide.

**Supplemental Table S2:** Predicted TOPs cleavage substrates including their DAP median |LFC| and the corresponding timepoint.

**Supplemental Table S3:** Scoring of previously found bioactive peptides with the Markov-Chain model.

**Supplemental Table S4:** Predicted bioactive peptides with significantly high Z-scores and differential abundance between *top1top2* and WT in the peptidome.

**Supplemental Table S5:** Intersection between predicted bioactive peptides and predicted TOP cleavage substrates.

**Supplemental Table S6:** Set of peptides selected for bioactivity screening.

**Supplemental Table S7:** Colony-forming units (CFU) (logarithmic scale) measured at 0 dpi and 3 dpi and normalized for plant weight to assess the impact of peptide treatments on plant response to PstAvrRpt2.

**Supplemental Data Set S1:** Quantified peptide abundances from LC-MS/MS data analysis. Quantified peptide abundances for Col-0 replicates are found in columns AK-AS. Quantified peptide abundances for *top1top2* replicates are found in columns AT-BB.

**Supplemental Data Set S2:** Statistical decision - genotype comparison.

**Supplemental Data Set S3:** Statistical decision - time series analysis.

**Supplemental Data Set S4:** GO enrichment analysis.

**Supplemental Data Set S5:** TOPs cleavage prediction results.

